# AIBP-CAV1-VEGFR3 axis dictates lymphatic cell fate and controls lymphangiogenesis

**DOI:** 10.1101/2020.10.16.342998

**Authors:** Xiaojie Yang, Jun-dae Kim, Qilin Gu, Qing Yan, Jonathan Astin, Philip S Crosier, Pengchun Yu, Stanley G Rockson, Longhou Fang

**Author notes:** Contributed equally (Xiaojie Yang, Jun-dae Kim). Prior lab members (Xiaojie Yang, Qilin Gu).

## Abstract

The lymphatics are essential for the maintenance of tissue fluid homeostasis. Accordingly, lymphatic dysfunction contributes to lymphedema. In development, lymphangiogenesis often requires lymphatic endothelial cell (LEC) lineage specification from the venous ECs and subsequent LEC proliferation and migration, all of which are regulated by the VEGFC/VEGFR3 signaling. Cholesterol is essential for proper cell functions and organ development, yet the molecular mechanism by which cholesterol metabolism controls lymphangiogenesis is unknown. We show that the secreted protein, ApoA1 binding protein (AIBP), dictates lymphatic vessel formation by accelerating cholesterol efflux. Loss of Aibp2, the human paralog in zebrafish, impairs LEC progenitor specification and impedes lymphangiogenesis. Mechanistically, we found that caveolin-1 (CAV-1) suppresses VEGFR3 activation in LECs, and that AIBP-regulated cholesterol efflux disrupts lipid rafts/caveolae and reduces CAV-1 bioavailability, which abolishes the CAV-1 inhibition of VEGFR3 signaling, thereby augmenting VEGFR3 activation and increasing lymphangiogenesis. Enhancement of cholesterol efflux with ApoA1 overexpression or inhibition of cholesterol biosynthesis using atorvastatin restores proper lymphangiogenesis in Aibp2 mutant zebrafish. Loss of Cav-1 increases LEC progenitor specification in zebrafish, and rescues lymphangiogenesis in Aibp2-deficient animals. Recombinant AIBP supplement confers profound LEC fate commitment in the mouse embryonic stem cells (mESC) to LEC differentiation model. Furthermore, enhancement of AIBP-CAV-1-VEGFR3 signaling axis promotes VEGFC-engaged adult lymphangiogenesis in mice. Consistent with these data, AIBP expression is reduced in the epidermis of human lymphedematous skin. These studies identify that AIBP-mediated cholesterol efflux is a critical contributor for lymphangiogenesis. Our studies will provide a new therapeutic avenue for the treatment of lymphatic dysfunctions.

**One Sentence Summary:** Our studies identify that AIBP-CAV-1-VEGFR3 axis enhances VEGFC-elicited lymphangiogenesis, which will guide a new therapeutic strategy for the treatment of lymphatic dysfunctions.

## Introduction

Lymphatic vasculature is essential for maintaining interstitial fluid homeostasis, dietary lipid transport and immune surveillance (*1–3*). The Lymphatic contribution is implicated in the pathogenesis of a variety of diseases, including lymphedema, tumor metastasis, cardiovascular disease, obesity, and diabetic mellitus (*4–11*). Therapeutic augmentation of lymphangiogenesis has been documented to improve lymphatic structure and function (*1–3, 7, 8*). Earlier studies show that the lymphatic system is derived from the embryonic cardinal vein (CV), where lymphatic endothelial cells (LEC) progenitors are specified and subsequently migrate and establish the lymphatic network (*1–3, 12, 13*). Recent studies indicate that a subset of lymphatics can be developed by non-canonical mechanisms (*14–19*). During murine development, the assembly of the lymphatic vascular network is initiated at approximately embryonic day 9.5 (E9.5) (*20*) under the control of VEGFC signaling through its cognate receptor VEGFR3 (*21, 22*). LEC fate commitment occurs through SOX18-induced expression of PROX-1, which, in concert with the orphan nuclear factor NR2F2 (also known as COUP-TFII), dictates LEC differentiation from the CV (*23, 24*). In addition, GATA2, HHEX, and PROX1 itself have recently been shown to upregulate PROX1 expression (*25–28*). The newly specified LECs subsequently bud from the CV and migrate (E10.0-E11.5) in a dorsolateral fashion to form lymph sacs and, thereby establish the entire lymphatic vessels. VEGFC-induced VEGFR3 signaling is the major driver of lymphangiogenesis in vertebrates. Interestingly, in addition to the regulation of LEC sprouting, VEGFR3 signaling also controls LEC progenitor homeostasis by maintaining PROX1 expression levels in a positive feedback loop (*21*). In zebrafish, Vegfc/Vegfr3 signaling increases Prox1^+^ LEC progenitors in the axis vasculature (*29, 30*). Mice deficient either in CCBE1, a critical matrix protein regulating VEGFC bioavailability, in VEGFC, or in VEGFR3 show fewer PROX1-positive LECs in the CV (*22, 31, 32*).

The developmental venous origin of the lymphatic endothelium and the requirement of VEGFC/VEGFR3 signaling for lymphangiogenesis are highly conserved from zebrafish to mice to humans (*20, 33*). In zebrafish, from 30 to 32 hours post fertilization (hpf), bipotential precursor cells in the posterior cardinal veins (PCV) express Prox1 and undergo a Vegfc/Vegfr3-dependent cell division to generate lymphatic and venous daughter cells. In response to the Vegfc cue, the daughter cells on the PCV progressively acquire the LEC fate at 36 hpf; among these, approximately half form parachordal LECs (PL) at the horizontal myoseptum by 48 hpf (*30, 34*). These PLs subsequently migrate ventrally and dorsally along the arterial intersegmental vessels (ISVs) at ~60 hpf (*34*), to form the thoracic duct (TD), intersegmental lymphatic vessels and dorsal longitudinal lymphatic vessels (DLLV) (*20*). Wnt5b has been reported to function upstream of Prox1 to regulate proper lymphatic specification (*15*). Nevertheless, the dynamics of LEC fate determination and the molecular mechanism of this process remain to be further understood.

We have demonstrated that the secreted protein AIBP limits angiogenesis in a non-cell autonomous fashion. Mechanistically, extracellular AIBP binds ECs, accelerates cholesterol efflux from ECs to high-density lipoprotein (HDL) and reduces lipid raft/caveola abundance, which in turn disrupts VEGFR2 signaling, thereby restricting angiogenesis (*35, 36*). Lymphatic vessels are structurally and functionally related to blood vessels. Many genes that are required for angiogenesis, such as VEGFR3, FGF, Dll4-Notch, angiopoietin-Tie2, Ephrin B2, and TGFβ family member ALK1, Epsin, and others, also function in lymphangiogenesis (*4, 37–47*). Thus, we sought to explore the role AIBP-regulated cholesterol metabolism in lymphangiogenesis. Our studies reveal a previously unidentified role of AIBP in lymphangiogenesis, in which AIBP augments VEGFC-engaged VEFGR3 signaling in a CAV-1-dependent fashion.

## Results

### Depletion of Aibp2 impairs lymphangiogenesis in zebrafish

We investigated lymphatic vessel formation in the recently generated Aibp2 knockout zebrafish (*48*). The *apoa1bp2^-/-^* zebrafish were crossbred with *fli1a:egfp* zebrafish that express EGFP in the both the lymphatic and blood ECs (*34, 49*), generating *apoa1bp2^-/-^; fli1a:egfp*. The TD is the first functional lymphatic vessel formed in the zebrafish trunk, situated between the dorsal aorta (DA) and PCV. At 5 days post fertilization (dpf), control siblings have a clearly demarcated TD running between the DA and PCV, whereas ~90% of *apoa1bp2^-/-^* embryos displayed ≤30% length of normal TD. Notably, ~80% of mutants showed a complete loss of this primary lymphatic vessel (Fig. 1A and B). Since the TD arises from the migration of PLs from the horizontal myoseptum, we assessed the development of PLs in the presence or absence of Aibp2. As illustrated in Fig. 1C and D at 48 hpf, the PL string was normally formed in control embryos, but was completely abolished in ~82% of *apoa1bp2^-/-^* animals and formed in a few segments (0-30%) in ~8% of animals. Similar developmental defects of TD and PLs were observed by morpholino antisense oligo (MO)-mediated *apoa1bp2* knockdown, which were restored to a large extent by co-injection of *apoa1bp2* mRNA (fig. S1A-C). Impaired TD formation can also be found in *apoa1bp2^-/-^; lyve1:Dsred* zebrafish (fig. S1D&E), which express DsRed in the lymphatic and venous ECs (*50*). Taken together, our data suggest that Aibp2 regulates lymphangiogenesis.

**Fig. 1.**
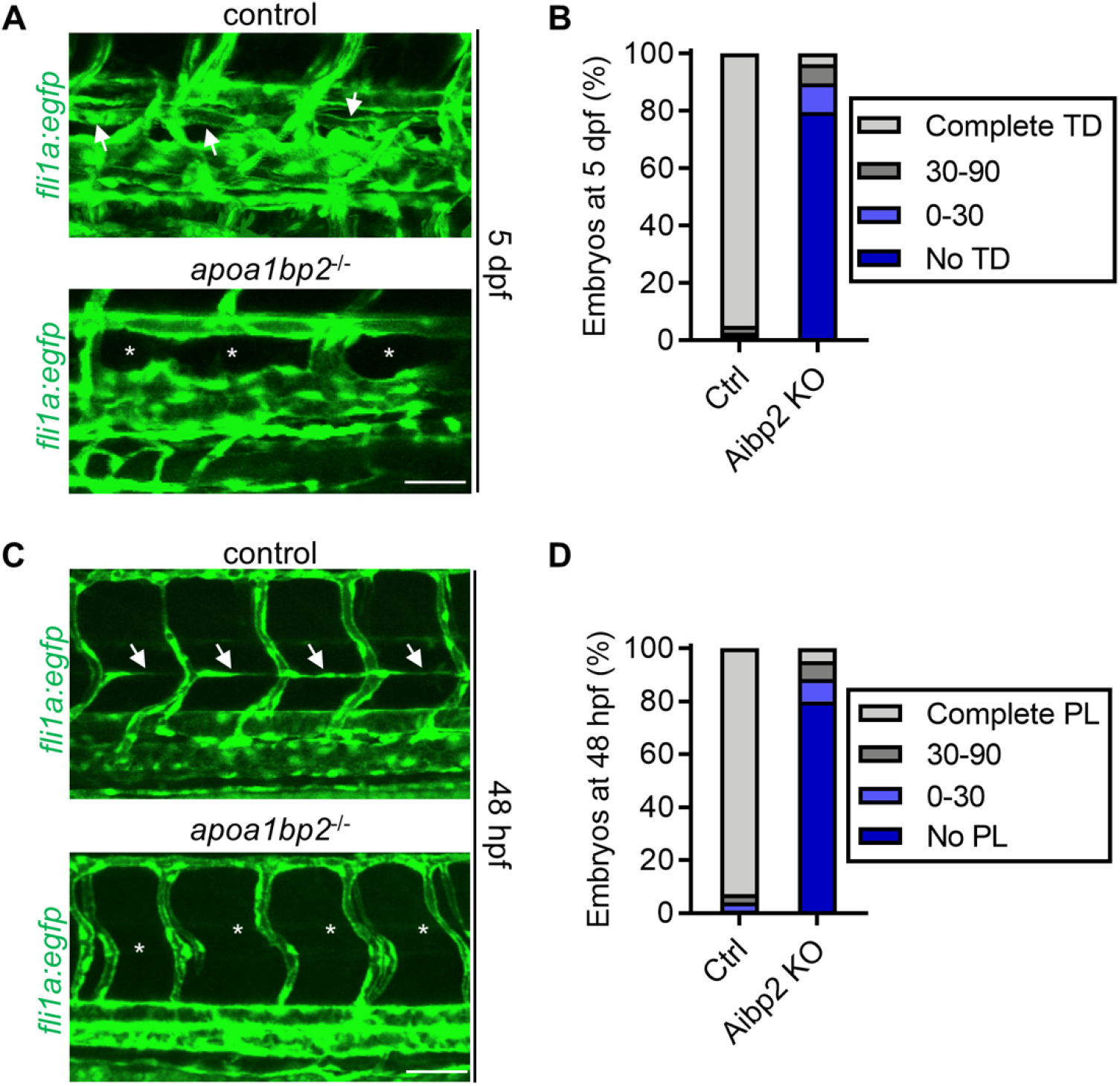
Effect of Aibp2 knockout on lymphatic vessel development. (**A**) Lymphatic developmental defects in *apoa1bp2^-/-^* zebrafish. Maxi-projection confocal images of TD formation in the *flila:egfp* and *apoa1bp2^-/-^; flila:egfp* null zebrafish at 5 dpf. Arrows show the TD, and stars denote absent TD. (**B**) Quantitative data of TD formation in A. n=15. (**C**) Loss of Aibp2 disrupts PL formation in *flila:egfp* zebrafish. Confocal images of PL in the horizontal myoseptum at 48 hpf. Arrows indicate PLs, and stars show absent PLs. (**D**) Quantitative data of the PL string in (C). n=20. Scale: 25 μm.

### AIBP enhances LEC lineage specification in zebrafish and in the mouse embryonic stem cells (mESCs) to LEC differentiation model

The defects in PL formation provoked us to explore whether Aibp2 has any effect on lymphatic progenitor specification by immunostaining of Prox1 at 36 hpf. Prox1 is the master transcription factor that dictates LEC fate specification, and is used as a readout of LEC identity (*27, 29, 30*). In control siblings, ~ 3 of Prox1-positive LEC progenitors per 6 segments were found in the PCV. By contrast, Aibp2 knockout markedly reduced the number of lymphatic precursors to ~1 (Fig. 2A and B), suggesting that Aibp2 regulates the development of lymphatic progenitors. We also analyzed the lymphatic vessel formation in embryos injected with *apoa1bp2* MO at 36 hpf. Reduced numbers of LEC progenitors in Aibp2 morphants corroborates the Aibp2 effect on lymphatic progenitor specification (fig. S2A and B).

**Fig. 2.**
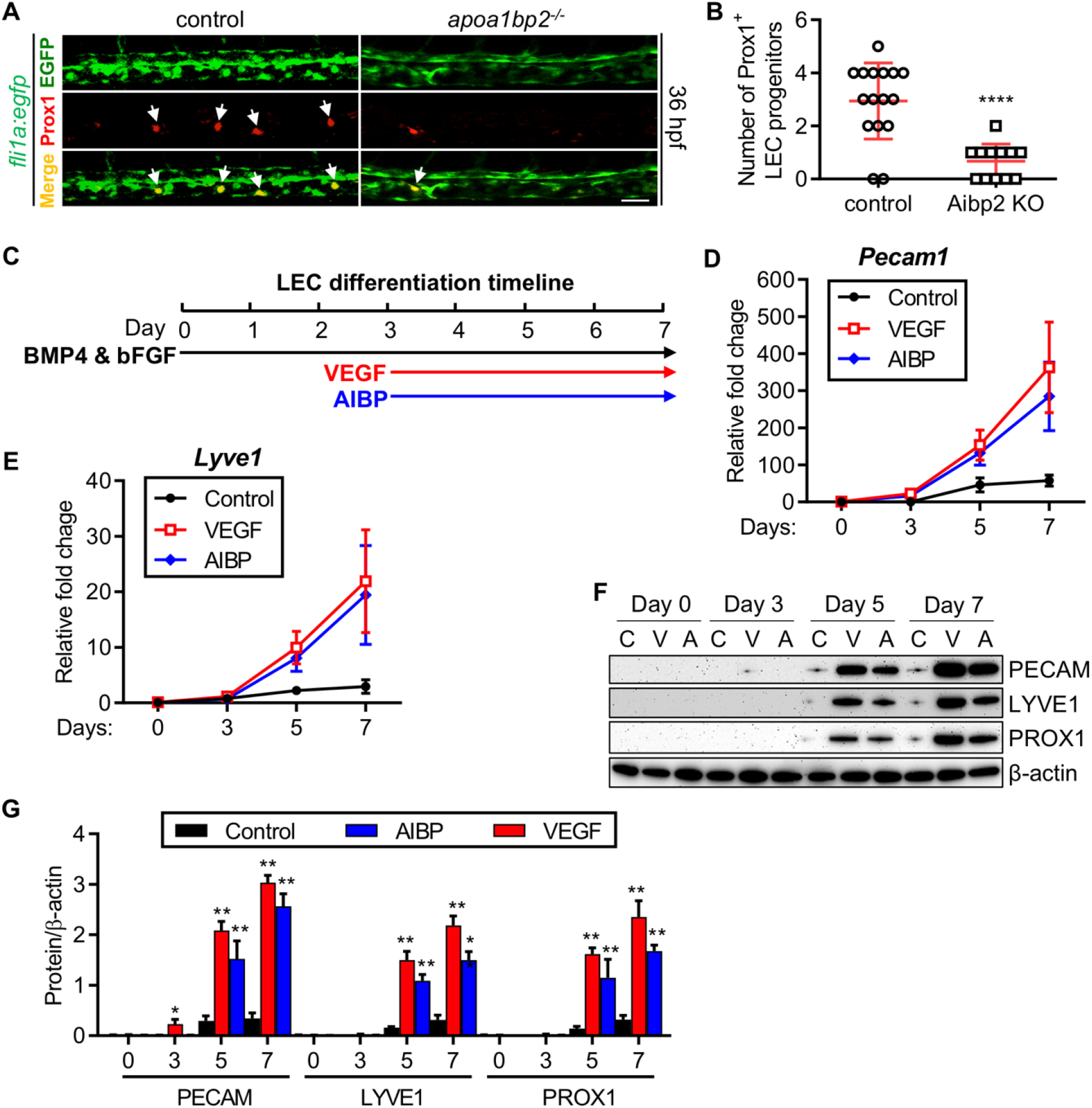
AIBP effect on LEC lineage commitment. (**A**) Impaired LEC specification in *apoa1bp2^-/-^; fli1a:egfp* zebrafish. Control and *apoa1bp2* knockout zebrafish were fixed with 4% PFA at 36 hpf, whole mount immunostaining was performed using Prox1 and EGFP antibodies, and images were captured using confocal microscopy. (**B**) Enumeration of LEC progenitors in panel A. Arrows show specified LEC progenitors. (**C**) Scheme illustration of mESC to LEC differentiation. (**D** to **G**). AIBP augments LEC differentiation from mESCs. The embryoid bodies (EBs) of mESCs were prepared and cultured in the EC differentiation medium containing 2 ng/ml BMP4 and 10 ng/ml bFGF for 3 days. Recombinant VEGFA (50 ng/ml) and VEGFC (50 ng/ml) in combination or AIBP (100 ng/ml) alone were supplemented at day 3 and kept in culture for additional 4 days. The resulting cells were harvested for qPCR analysis of endothelial cell (EC) marker *Pecam1* expression (**D**) and LEC-associated *Lyve1* expression (**E**) at the indicated time points. Western blot analyses of PECAM, LYVE1 and PROX1 expression (**F**) and the quantitative data (**G**). C: control; V: VEGFA + VEGFC; A: AIBP. *, p<0.05; **, p<0.001; ****, p<0.0001. Scale: 25 μm.

We next determined whether the role of AIBP in lymphatic progenitor specification is evolutionarily conserved. Mouse embryoid bodies (mEBs) prepared from mESCs can be differentiated into the derivatives of ectodermal, mesodermal, and endodermal tissues and recapitulate certain developmental processes (*51*); the LECs emerge from the mesoderm. We prepared mEBs and assessed LEC generation in the differentiation medium containing BMP4 and bFGF followed by additional supplement on day 3 through day 7 with recombinant VEGFA and VEGFC in combination or with AIBP alone (Fig. 2C). From day 0 onwards during differentiation, LECs were identified by the expression of PECAM, LYVE1 and PROX1, as previously reported (*52*). Compared with control cells, VEGFA and VEGFC co-treatment increased both the mRNA and protein expression levels of these LEC-associated markers, LYVE1 and PROX1, at day 5 and day 7 of differentiation (Fig. 2D-G). Surprisingly, AIBP incubation strikingly increased LEC progenitor specification as evidenced by robust expression of the LEC markers (Fig. 2D-G), comparable to the effect of VEGFA/C co-administration. A greater percentage of PECAM^+^/LYVE1^+^ LEC population was detected by FACS analysis at day 7 of LEC differentiation in the presence of AIBP or VEGFA/C (fig. S3A and B). These results demonstrate that AIBP regulation of LEC fate is conserved across evolution.

### Aibp2-mediated cholesterol efflux is essential for proper lymphangiogenesis

Our previous studies document that AIBP accelerates cholesterol efflux from ECs (*36*). Thus, we first measured free cholesterol levels in *apoa1bp2^-/-^* embryos. Indeed, Aibp2 depletion significantly increases free cholesterol content in *apoa1bp2^-/-^* embryos (Fig. 3A). We then treated *apoa1bp2* animals with atorvastatin, a cholesterol-lowering drug, and assessed the attendant effect on lymphangiogenesis. We found that ~93% of *apoa1bp2^-/-^* embryos displayed ≤50% TD formation, and within this population, ~80% of the embryos lacked the TD. In contrast, atorvastatin treatment substantially reduced the percentage of *apoa1bp2^-/-^* mutants with incomplete TD development, suggesting that atorvastatin-mediated reduction of cell cholesterol content rescues lymphatic vessel formation (Fig. 3B and C). In addition, we overexpressed apoA-I, the major protein constituent of HDL, in *apoa1bp2* knockout zebrafish. Similarly, enforced human apoA-I expression corrected TD formation in *apoa1bp2* knockout animals, as evidenced by ~76% of *apoa1bp2^-/-^* zebrafish receiving *APOA1* mRNA show rescued TD development (Fig. 3D and E). These results indicate that Aibp2 regulates lymphatic vessel formation through cholesterol metabolism.

**Fig. 3.**
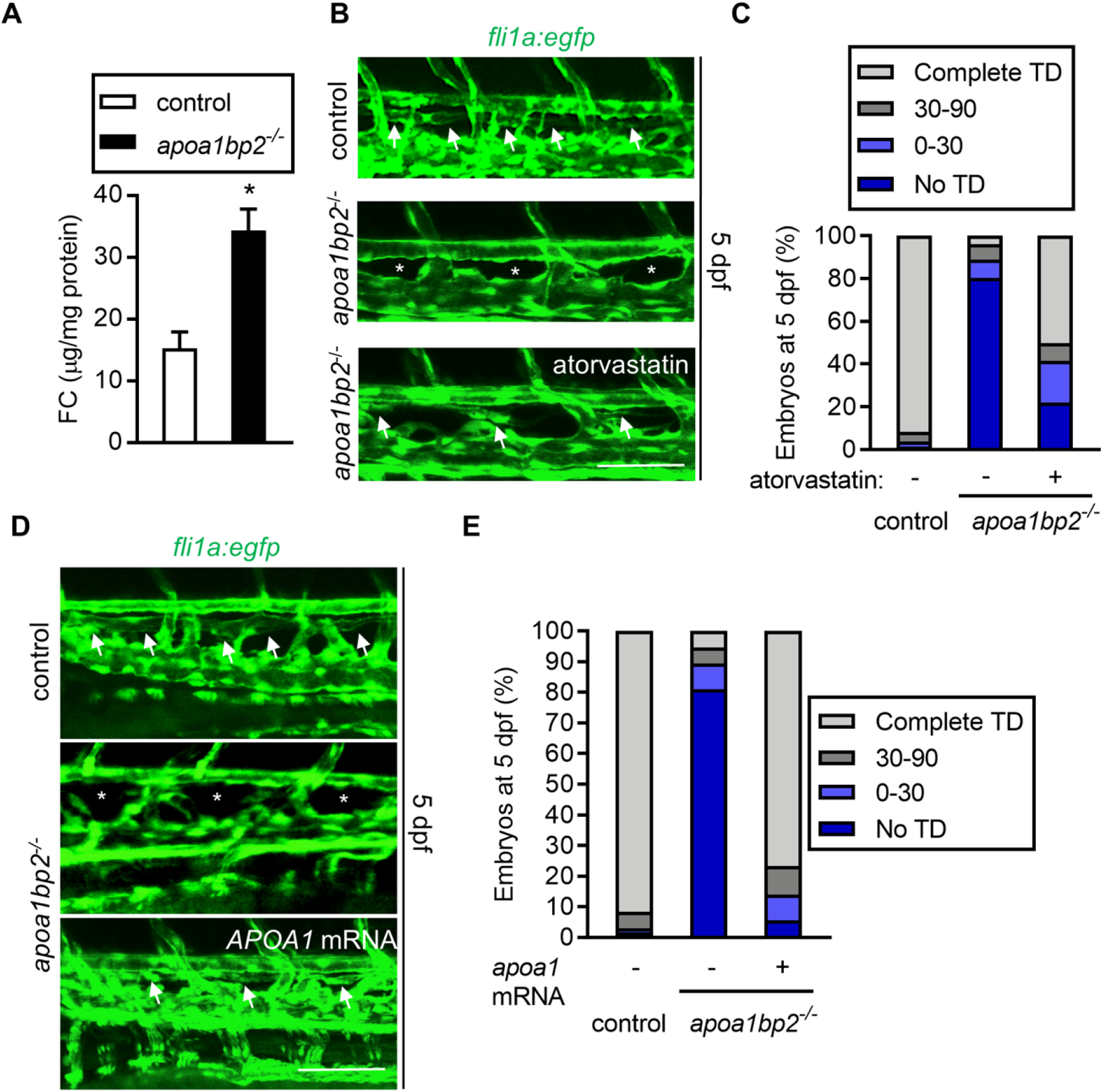
Effect of cholesterol reduction on lymphatic vessel development. (**A**) Free cholesterol (FC) content in control and Aibp2 null zebrafish. The 5 dpf zebrafish trunk of control or Aibp2 null zebrafish were dissected, total lipids extracted, and free cholesterol levels measured. The FC content were normalized to the protein levels. Fifty embryos were pooled for each measurement. Mean±SE; *, p<0.05. (**B**) Inhibition of cholesterol synthesis rescues TD formation in Aibp2 null animals. TD formation in control zebrafish, Aibp2 null zebrafish treated with 1 μM atorvastatin or control vehicle ethanol at 5 dpf. (**C**) Quantitative data of TD formation in B. n=20. (**D**) apoA-I-mediated cholesterol efflux restores TD formation. Confocal images of TD formation in control zebrafish, Aibp2 null zebrafish, or Aibp2 knockout zebrafish with 100 ng human *APOA1* mRNA overexpression. (**E**) Quantitative data of TD formation in D. Arrows show TD, and stars denote absent TD. n=18. Scale: 25 μm. *, p<0.05.

### AIBP-mediated cholesterol efflux disrupts caveolae and enhances VEGFC/VEGFR3 signaling in human LECs

CAV-1, a structural protein organizing the formation of caveolae, binds cholesterol directly, thereby forming cholesterol-rich microdomains, which provide a platform to facilitate membrane-anchored receptor signaling (*53, 54*). CAV-1 ablation in mice eliminates caveolae (*55, 56*). Both the expression of CAV-1 and the formation of caveolae are dependent on cellular cholesterol levels. In many cases, CAV-1 deficiency alters plasma membrane receptor signaling competence through disruption of caveolae (*57–59*).

Following LEC specification from the venous ECs, VEGFR3 signaling is required for the maintenance of the LEC progenitor fate, as well as LEC proliferation and migration (*30*). CAV-1 in caveolae was reported to repress VEGFR3 activity in the ECs (*60*). We postulate that AIBP-mediated cholesterol efflux regulates lymphangiogenesis through CAV-1-dependent VEGFC/VEGFR3 signaling. As shown in Fig. 4A and B, compared with VEGFC treatment in human LECs (hLECs), methyl beta cyclodextrin (MβCD)-mediated cholesterol depletion profoundly increased VEGFC-induced VEGFR3 activation, suggesting that caveolae abundance and associated CAV-1 bioavailability inhibit VEGFR3 signaling. Given that AIBP-mediated cholesterol efflux disrupts caveolae, we tested whether AIBP exerts its effect on CAV-1-regulated VEGFC/VEGFR3 signaling via modulation of caveolae. We performed detergent-free, discontinuous gradient ultracentrifugation analysis of hLECs, which were pre-incubated with control media, AIBP, HDL_3_, or their combination. As anticipated, VEGFR3 was present in the caveolar fractions that were positive for CAV-1 (Fig. 4C). AIBP and HDL_3_ treatment in concert, which disrupts caveolae, decreased CAV-1 localization in the caveolar fraction and induced VEGFR3 redistribution from the caveolar to the non-caveolar domains (Fig. 4C). To further investigate the effect of AIBP-mediated cholesterol efflux on VEGFR3 activation, we treated hLECs with AIBP, HDL_3_, or their combination, followed by stimulating cells with VEGFC. We found that AIBP or HDL_3_ alone did not significantly affect VEGFC-induced activation of AKT and ERK, the two downstream effector kinases. However, AIBP and HDL_3_ in combination, similarly to the treatment with MβCD (Fig. 4B), facilitated VEGFC-induced VEGFR3 activation as shown by markedly increased phosphorylation of AKT and ERK (Fig. 4D-F). These results suggest that AIBP-mediated cholesterol efflux disrupts plasma membrane caveolae, thereby facilitating lymphangiogenesis by augmenting VEGFR3 activation.

**Fig. 4.**
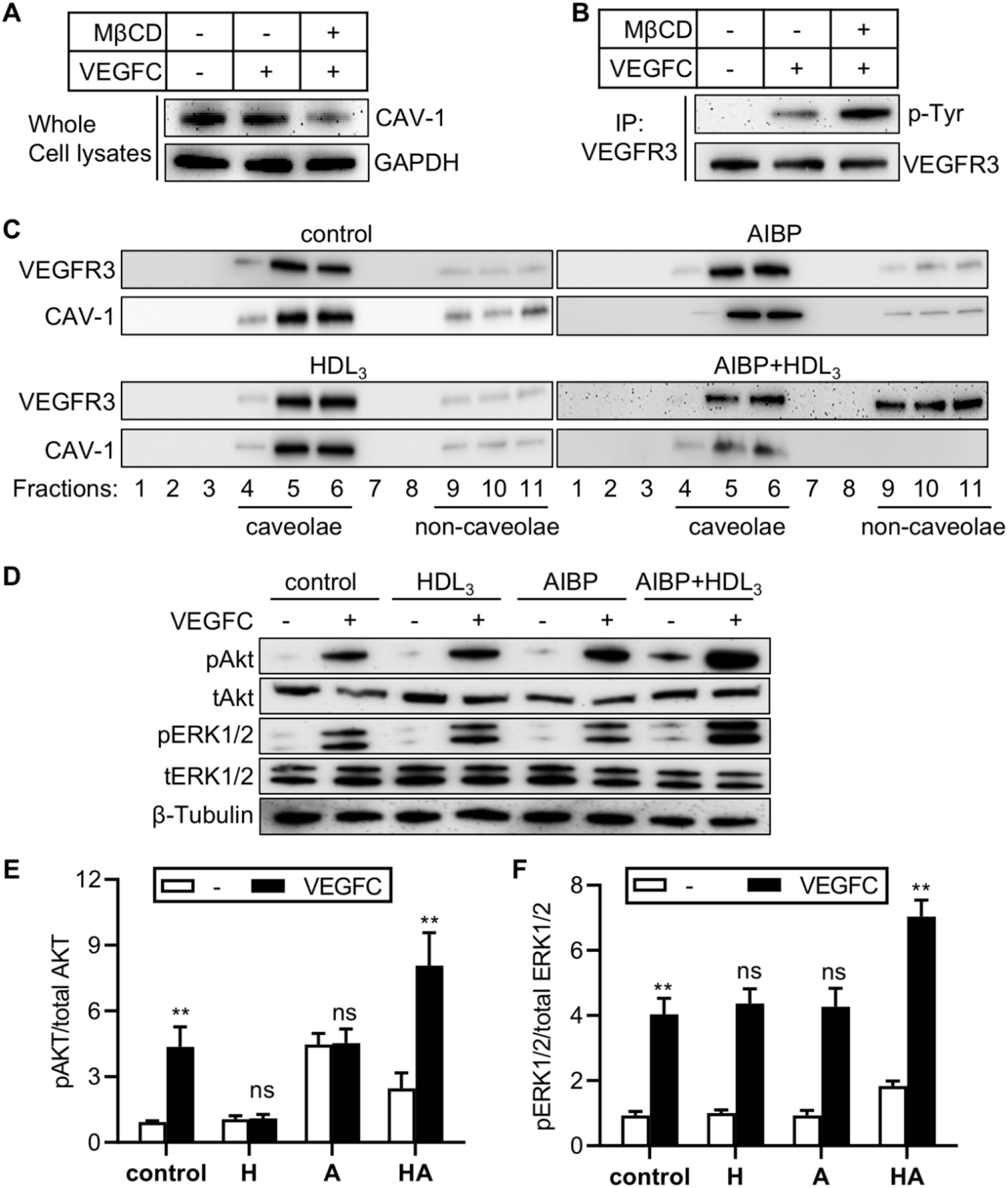
Effect of cholesterol efflux on VEGFR3 signaling. (**A** and **B**). Cholesterol removal potentiates VEGFR3 signaling. (**A**) hLECs were serum-starved, and treated with 10 mM MβCD for 30 min, and cells were further stimulated with 100 ng/ml VEGFC. The resulting cells were lysed and blotted with Cav-1 or GAPDH antibodies. (**B**) hLECs were treated as in (**A**). and cell lysates were immunoprecipitated using VEGFR3 antibody. Immunoblotting was performed using anti-phosphotyrosine (4G10) and VEGFR3 antibodies. (**C**) AIBP-mediated cholesterol efflux disrupts caveolae and reduces CAV-1 levels in the caveolar fractions. hLECs were treated with recombinant 200 ng/ml AIBP, 100 μg/ml HDL_3_, or both in serum-free EBM2 for 6 hours, and the cells were subjected to sucrose-mediated ultracentrifugation. The resulting fractions were collected for Western blot analysis as indicated. (**D**) AIBP-mediated cholesterol efflux increases VEGFR3 signaling. hLECs were serum-starved and treated as in (**C**), and further stimulated with 100 ng/ml VEGFC. The resulting cells were lyzed and immunoblotted as indicated. (**E** and **F**), Quantitative data of AKT activation (**E**) and ERK activation (**F**). Mean±SD, n=3 independent repeats. *, p<0.05; **, p<0.01; ns: not significant.

### Disrupted CAV-1 and VEGFR3 interaction augments VEGFR3 signaling in hLECs

Whereas Cav-1 is a core component of caveolae, it is also a scaffolding protein that provides a spatially restricted platform for proper signaling of cell surface receptors (*61*). Cav-1 contains a signaling motif that interacts with a variety of membrane receptors and modulates their activities (*62*). We found that VEGFR3 contains such a conserved motif (Fig. 5A), implying possible regulation of its signaling by CAV-1. We thus generated a VEGFR3 mutant VEGFR3^AAA^ that lacks the three conserved amino acids in the hypothetical CAV-1 binding domain. We overexpressed EGFP, VEGFR3, or VEGFR3^AAA^ in hLECs using lentivirus-mediated gene transfer and examined Cav-1/VEGFR3 interaction. As illustrated in Fig. 5B, wild-type VEGFR3 but not VEGFR3^AAA^ binds CAV-1 in hLECs.

**Fig. 5.**
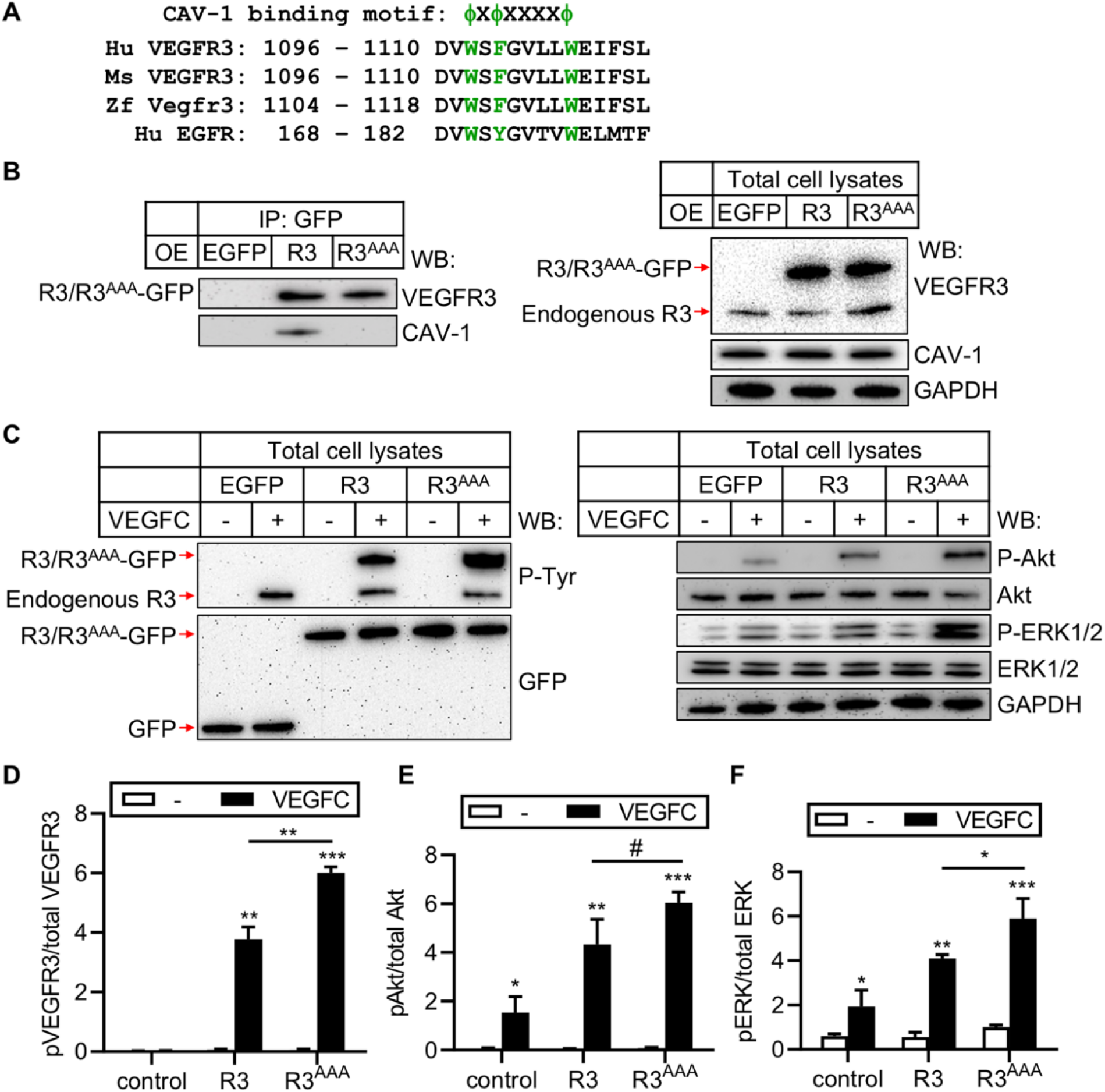
Effect of CAV-1 on VEGFR3 signaling. (**A**) Conserved CAV-1 binding site on VEGFR3 in human (Hu), mouse (Ms), and zebrafish (Zf). (**B**) VEGFR3^AAA^ loses its binding to CAV-1. hLECs were transfected with control EGFP (Ctrl), VEGFR3-EGFP (R3), or VEGFR3^AAA^-EGFP (R3^AAA^) using lentivirus-mediated gene transfer. After 72 hours, the resulting cells were lyzed and immunoprecipitated with EGFP antibody coupled to the magnetic Dynabeads and immunoblotted using VEGFR3 and CAV-1 antibodies. Immunoblotting of overexpressed VEGFR3-EGFP and VEGFR3^AAA^-EGFP (R3/R3^AAA^-EGFP) were detected using VEGFR3 antibody. The input lysates were shown on the right. (**C**) VEGFR3^AAA^ increases VEGFR3 signaling. hLECs were transduced as in (**B**), and the resulting cells were serum starved and treated with 100 ng/ml VEGFC for 20 min, cells were then lysed and immunoblotted as indicated. Quantitative data of VEGFR3 activation (**D**), AKT activation (**E**), and ERK activation (**F**) were shown. n=3 independent repeats. *, p<0.05; **, p<0.01; ***, p<0.001.

To test the effect of the VEGFR3^AAA^ mutant on VEGFR3 signaling, we ectopically expressed EGFP, VEGFR3 and VEGFR3^AAA^ in hLECs using lentivirus-mediated transfection, followed by stimulation with VEGFC, and assessed VEGFR3 activation. The results show that the triple Ala point mutations, which disrupt its interaction with CAV-1 (Fig. 5B), results in potent VEGFC-induced VEGFR3 phosphorylation as well as AKT and ERK activation compared to wild-type VEGFR3 (Fig. 5C-F). Taken together, our data suggest that AIBP-mediated cholesterol efflux disrupts plasma membrane caveolae and mitigates CAV-1-mediated inhibition of VEGFR3 signaling in LEC.

### Cav-1 regulates LEC progenitor development

To explore the role of Cav-1 in lymphatic vessel formation, we generated *cav-1^-/-^* mutant zebrafish (fig. S4A-C), in which Western blot analysis revealed the absence of Cav-1 protein (Fig. S4D). Morphologically, Cav-1 null zebrafish are comparable to control animals from 6 hpf to 5 dpf (fig. S4E). Interestingly, Cav-1 knockout zebrafish showed greater numbers of Prox1 labeled LEC progenitors at 36 hpf (Fig. 6A and B). However, TD formation appeared normal in these animals (Fig. 6C and D). Cav-1 depletion or overexpression resulted in profoundly increased or reduced expression of lymphatics-associated genes at 96 hpf, respectively (fig. S5A and B). Thus, our data show that Cav-1 suppresses lymphatic progenitor development.

**Fig. 6.**
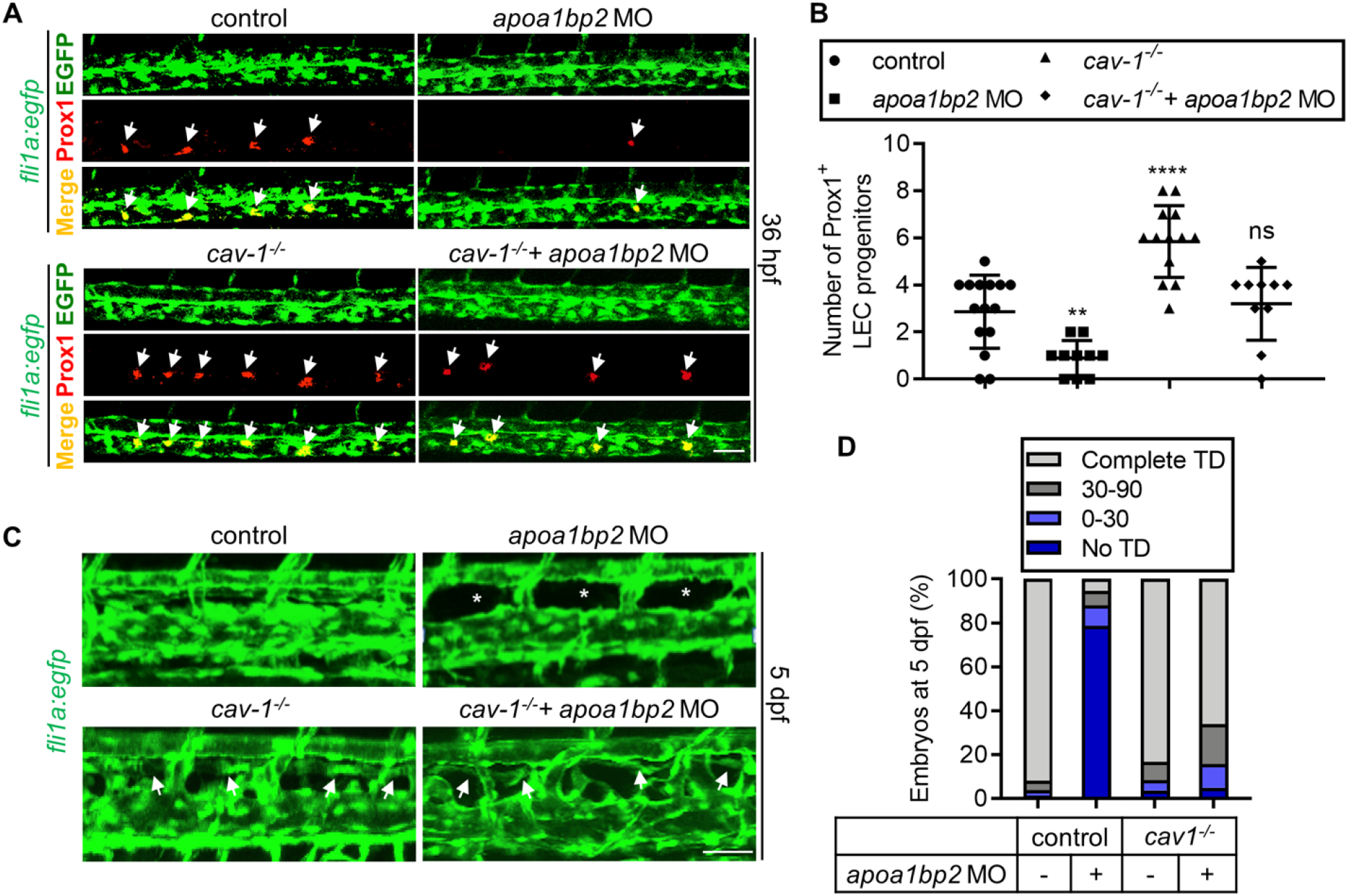
Effect of CAV-1 on Aibp2-regulated LEC specification. (**A**) Cav-1 deficiency rescues LEC specification in Aibp2 knockdown animals. Max projection confocal images of control, Aibp2-deficient, Cav-1 null, or Cav-1/Aibp2 double deficient zebrafish. The animals were PFA fixed at 36 hpf and immunostained using Prox1 and EGFP antibodies. Arrows indicate LEC progenitors. (**B**) Quantitative data of LEC progenitors in (**A**). (**C**) Cav-1 deficiency corrects lymphatic defects in Aibp2 knockdown animals. Confocal imaging of TD formation in the indicated genetically modified *fli1a:egfp* zebrafish at 5 dpf. Arrows show TD, and stars denote absent TD. (**D**) Quantitative data of TD formation in (**C**). **, p<0.01; ****, p<0.0001; ns: not significant, n=20. Scale, Scale: 25 μm.

### Loss of Cav-1 rescues defective lymphangiogenesis in Aibp2-deficient zebrafish

We have shown that Aibp2 knockout increases free cholesterol content (Fig. 3A), which presumably increases caveola content, as cholesterol is enriched in the caveolae (*63*). Given that CAV-1-organized caveolae negatively regulate VEGFR3 signaling (*60*), we examined the effect of Cav-1 deletion on lymphangiogenesis in Aibp2-deficient animals. As expected, absence of Cav-1 rescued impaired lymphatic progenitor development in Aibp2 knockdown animals as revealed by recovered Prox1 staining (Fig. 6A and B). Consistent with this rescue effect, defective TD formation was restored in the Cav-1- and Aibp2-double deficient animals (Fig. 6C and D). The data suggest that Cav-1 functions downstream of Aibp2-mediated cholesterol efflux.

### AIBP-mediated cholesterol efflux promotes adult lymphangiogenesis in mice

Since Cav-1 depletion rescues the TD development in Aibp2 morphants, we hypothesized that Aibp2, by alleviating Cav-1 repression of Vegfr3, facilitates lymphangiogenesis. We further investigated this mechanism in a murine model to determine whether this mechanism is evolutionarily conserved. Indeed, loss of CAV-1 (*55*) significantly augmented tail lymphangiogenesis in P3 neonatal mice as revealed by LYVE-1 antibody staining of lymphatic vessels in the similar anatomic location (fig. S6A and B). The corneas of adult mice lack lymphatic vasculature (*64, 65*), thereby providing a widely used model to study injury or growth factor-induced lymphangiogenesis (Fig. 7A). To explore the role of cholesterol efflux in adult lymphangiogenesis, we implanted VEGFC-containing pellets, in the presence or absence of AIBP and the core protein component of HDL – apoA-I or the CAV-1 blocking peptide (CAV-1 scaffolding domain, CSD) – into the corneas of B6 mice. We used apoA-I in this assay due to technical difficulties in the preparation of HDL pellets. As expected, VEGFC pellet implantation elicited robust lymphangiogenesis (Fig. 7B). Remarkably, co-administration with AIBP and cholesterol acceptor apoA-I further increased lymphangiogenesis (Fig. 7B and C). Similarly, supplement of CSD augmented lymphatic vessel formation (Fig. 7B and C). However, no pro-lymphangiogenic effect was found with AIBP/apoA-I or CSD treatment in the absence of VEGFC (fig. S7). Thus, consistent with our studies in zebrafish, AIBP-mediated cholesterol efflux increases lymphangiogenesis in neonatal and adult mice.

**Fig. 7.**
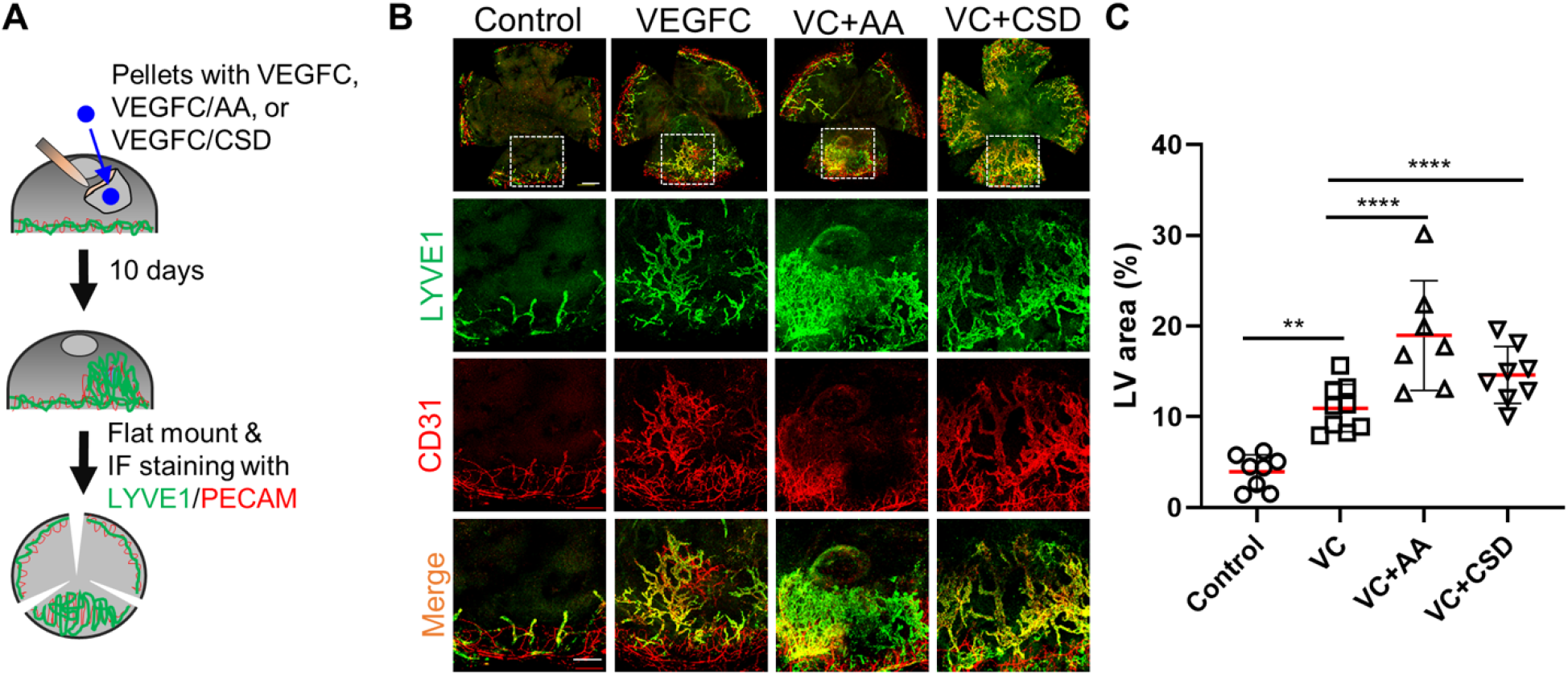
AIBP effect on corneal lymphangiogenesis and LEC specification. (**A**) Illustration of murine cornea lymphangiogenesis assay. (**B**) Representative images of murine corneal lymphangiogenesis by implantation of pellets containing the indicated recombinant proteins and immunostained using LYVE1 & CD31 antibodies. Enlarged images of the boxed regions (scale bar, 1000 μm) are shown in the lower panels (scale bar, 500 μm). (**C**) Quantification of LYVE1+ lymphatic vessel area per cornea. Ctrl: control; VC: VEGFC; AA: AIBP+ApoA-I; CSD: CAV-1 scaffolding domain peptide. **, p<0.01; ****, p<0.0001.

### Reduced AIBP expression in the affected epidermis of lymphedema patients

To probe the association of AIBP content with lymphatic vessel dysfunction, we examined AIBP expression in human skin specimens derived from 7 subjects. We obtained paired cutaneous normal and lymphedematous biopsy specimens from each of these individuals. Immunohistochemistry staining of AIBP showed that it is robustly expressed in the epidermis and the sweat glands (Fig. 8A). Interestingly, in human lymphedema, AIBP expression is mildly but significantly reduced in the epidermis but not in the dermis or sweat glands, when compared to the paired normal cutaneous specimens (Fig. 8A-D). The comparable expression in the dermis and sweat gland excludes the possibility that tissue swelling dilutes AIBP content in the epidermis of lymphedema biopsy samples. The observed abnormality in the lymphedematous epidermis suggests that reduced AIBP expression is associated with impairment of cutaneous lymphatic function.

**Fig. 8.**
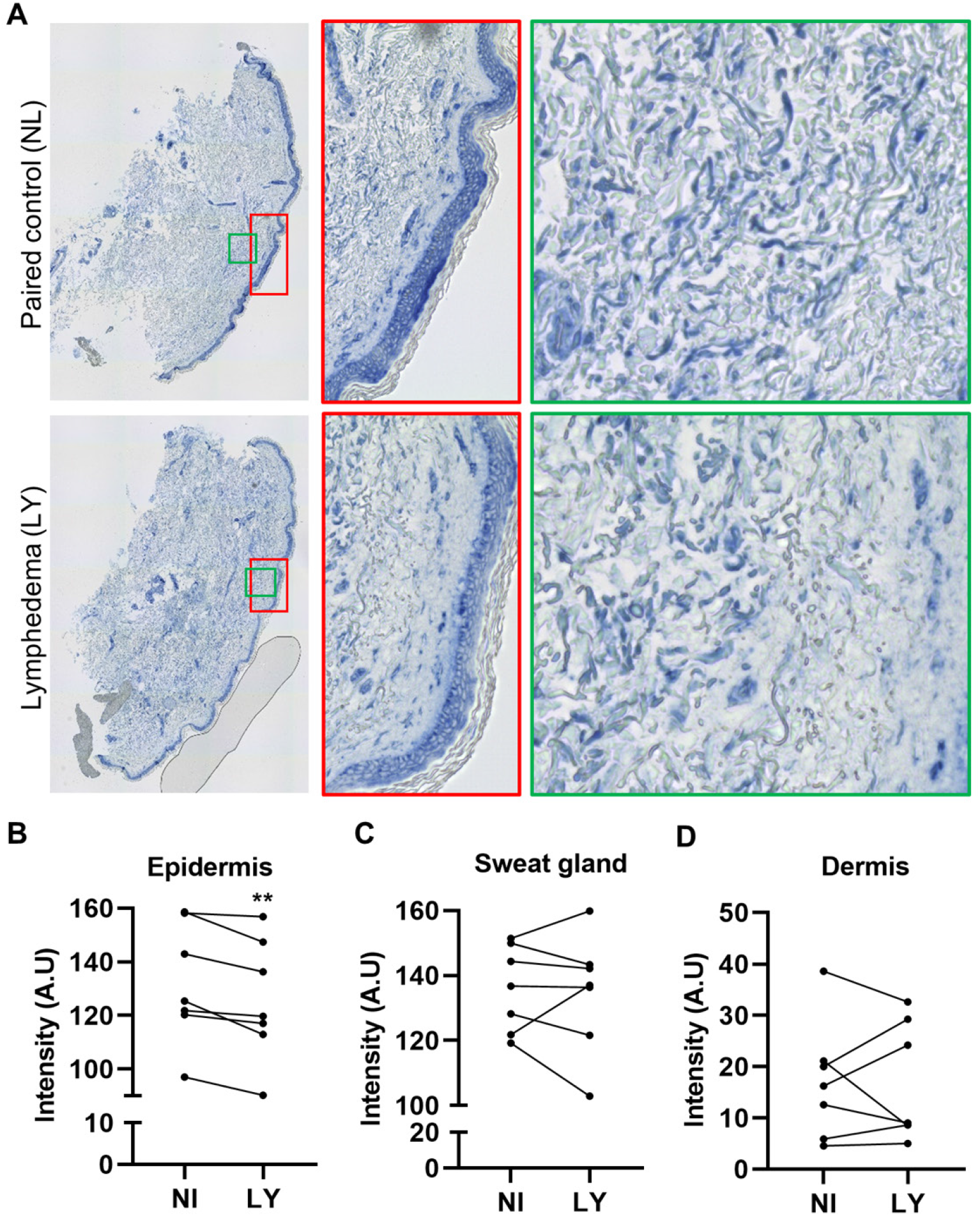
AIBP expression in cutaneous lymphedema. (**A**) Immunohistochemistry analysis of AIBP expression. Paired normal and lymphedema cutaneous biopsies (n=7 patients) were fixed with formalin and the 5 μm paraffin sections were prepared. The sections were deparaffinized and immunostained using our non-commercial AIBP antibody. (**B** to **D**), The AIBP signal in the epidermis (**B**), sweat glands (**C**), and dermis (**D**) and were quantified using ImageJ. The enlarged areas of green and yellow boxes are shown on the right. The images were taken separately and stitched together using the EVOS microscope. NL: normal leg; LY: lymphedema. **p<0.01.

## Discussion

In this paper, our studies reveal a novel molecular mechanism underlying lymphatic development, in which AIBP-mediated cholesterol efflux disrupts caveolae and, consequently, reduces CAV-1 bioavailability, thereby mitigating the CAV-1 inhibition of VEGFR3 activation and augmenting lymphatic fate commitment and lymphangiogenesis. Our mechanistic study of AIBP-regulated lymphangiogenesis will faciliate the development of new therapies for the diseases associated with lymphatic dysfunction.

### Opposing roles of AIBP in angiogenesis and lymphangiogenesis

Our previous studies have shown that AIBP-mediated cholesterol efflux disrupts lipid rafts/caveolae, which results in restricted angiogenesis by two mechanisms: 1) it impedes rafts/caveolae anchored proangiogenic VEGFR2 clustering, endocytosis and signaling, thereby restricting angiogenesis (*36*), and 2) it translocates γ-secretase from lipid rafts/caveolae to the non-raft/caveolar domain, facilitating the cleavage of NOTCH1 receptor and augmenting anti-angiogenic NOTCH1 signaling (*35*). Recently, we demonstrated that AIBP instructs hematopoietic stem and progenitor cell specification in development through activation of SREBP2, which in turn transactivates NOTCH1 for hematopoiesis (*48*). Here, we show that AIBP increases lymphangiogenesis and instructs LEC progenitor specification. At the molecular level, AIBP-mediated cholesterol efflux attenuates the inhibitory effect of CAV-1 on VEGFR3 signaling. Thus, disruption of CAV-1 and VEGFR3 binding augments VEGFR3 signaling and LEC progenitor development and lymphangiogenesis. Although LECs are derived from blood ECs (BEC), their functions and molecular identifications differ. The fundamental differences between BECs and LECs may contribute to the disparate effects of AIBP on angiogenesis and lymphangiogenesis.

It is not without precedent that one protein may have the opposing effects on angiogenesis and lymphangiogenesis. For instance, RhoB ablation augments lymphangiogenesis but impairs angiogenesis in the cutaneous wound healing model (*66*). Although Notch signaling limits angiogenesis (*67–69*), Notch has been reported to inhibit or augment lymphangiogenesis in a context-dependent fashion (*37, 38, 70, 71*).

### AIBP-mediated CAV-1 dependent regulation of VEGFR3 signaling and lymphangiogenesis

Lymphatic vessel formation requires LEC fate commitment and subsequent sprouting. The LEC fate acquisition, LEC sprouting from the embryonic CV, and dorsolateral migration under the control of VEGFC/VEGFR3/PROX1 signaling axis is highly conserved (*26*). Restricted expression of PROX1 protein dictates LEC identity, which is critical for future LECs to migrate out of the PCV (*27*). AIBP is evolutionarily conserved from zebrafish to human. Zebrafish have two *apoa1bp* genes that encode Aibp1 and Aibp2 proteins, and Aibp2 demonstrates functional properties resembling those of human AIBP (*36*). Although the degree of phenotypic severity varies, most of the Aibp2 mutants show a partial or complete loss of PLs in the horizontal myoseptum region and, thus, lacking large portion or all of the TD arising from PLs. Furthermore, these defects are likely attributed to a failure of LEC lineage commitment, as Aibp2 loss-of-function impairs lymphatic precursor specification (Fig. 2A and B).

A positive feedback loop between VEGFR3 and PROX1 is necessary to maintain the identity of LEC progenitors in mice and zebrafish (*21, 30*). CAV-1 has been reported to inhibit VEGFR3 signaling in ECs (*60*). Our studies show that Cav-1 knockdown or overexpression in zebrafish increases or decreases expression of lymphatic genes, respectively. Moreover, loss of Cav-1 strongly enhances lymphatic progenitor specification and rescues impaired lymphangiogenesis. Despite increased number of LEC progenitors, the TD formation appears normal in Cav-1 null zebrafish (Fig. 6C and D). Indeed, there are considerable variations in the number of LECs required to form a complete TD, as previously reported (*30*). CAV-1 deficiency augments lymphangiogenesis in mouse development and in adult cornea with VEGFC stimulation (Fig. 7A-C and fig. S6). In hLECs, VEGFR3 is present in caveolae and contains a binding motif for CAV-1 (Fig. 5A). The disruption of the CAV-1 and VEGFR3 interaction increases VEGFR3 activation in response to VEGFC treatment (Fig. 5C-F). These results suggest that AIBP-mediated cholesterol efflux facilitates lymphangiogenesis, which is achieved through abolishing the CAV-1-mediated inhibition of VEGFR3 signaling.

### AIBP effect on ERK signaling

ERK activation controls LEC fate specification and LEC sprouting (*29, 30, 72, 73*). A modest but sustained ERK activation in mice, resulting from the expression of a RAF mutant, results in lymphatic hyperplasia (*73*). Expression of the Vegfr3 mutant that abolishes Erk activation in zebrafish impairs development of Prox1^+^ LECs (*29*). Our data indicate that AIBP enhances VEGFC-mediated ERK activation in hLECs (Fig. 4D-F). Conversely, Aibp2 knockdown in zebrafish abrogates Erk1 activation while robustly facilitating Akt activation (*36*). Notably, VEGFR3 homodimer activates ERK while VEGFR2/VEGR3 heterodimer activates AKT (*74*). Our studies are consistent with the dominant role of ERK in lymphatic vessel development.

### Conserved role of AIBP in the regulation of LEC fate and lymphangiogenesis

In the mESCs to the LEC differentiation model, we found that recombinant AIBP incubation documents a remarkable effect on the LEC specification arising from mesodermal origin. Given that AIBP alone shows no observable effect on VEGFR3 phosphorylation, this AIBP effect on LEC fate commitment is likely due to a synergistic effect of AIBP and increased VEGFR signaling in the induced mesoderm (*75, 76*). Although AIBP is unlikely to initiate a ligand-independent signaling, it fine-tunes VEGFC/VEGFR3/ERK signaling.

Earlier studies suggest that LECs emerge from the venous origin (*2, 14, 15*). However, other mechanisms of lymphatic vessel formation have been reported recently, such as in the zebrafish facial lymphatics (*17*), murine dermal and cardiac lymphatic vessel formation (*18, 77*), and zebrafish brain (*16, 19*). The Aibp2 effect on lymphatic vessel formation is blunted at later developmental stages in zebrafish, and we also did not detect a significant difference in tail lymphangiogenesis between AIBP knockout neonatal mice and control littermates (data not shown), which could be due to the rescue effect of non-venous origin derived lymphangiogenesis.

It often takes years to develop secondary lymphedema, an acquired damage to the lymphatic system occasioned by surgical disruption, radiotherapy, or infection (*9, 78*). Genetic studies suggest that small changes in an array of lymphangiogenic genes, in the absence of a single dominant mutation, may precipitate secondary lymphedema (*79*). Our AIBP staining of paired human normal/lymphedema cutaneous biopsy specimens revealed that AIBP expression is significantly reduced in the epidermis in the settings of lymphedema, when compared to the paired, normal counterpart. Whereas it is a mild change, this may be indicative of a general attenuation of lymphangiogenic competence in the lymphedematous tissues.

## Materials and Methods

### Generation of Cav-1 knockout zebrafish using CRISPR/Cas9

The CRISPR/Cas9-mediated gene knockout technique was performed as previously described (*80*). Briefly, the target sequence of caveolin 1 (Cav-1, NCBI Gene ID: 323695) was 5’-GACGTGATCGCCGAGCCTGC-3’. The guide-RNA (gRNA) sequence was subcloned into pT7-gRNA vector and gRNA was synthesized using HiScribeTM T7 Quick High Yield RNA Synthesis (New England Biolabs). The pCS2-nCas9n was linearized using NotI and Cas9 mRNA was synthesized using mMESSAGE mMACHINE SP6 Kit (Invitrogen). Twenty-five ng/□l gRNA and 200 ng/μl Cas9 mRNA were mixed and injected into one-cell stage wild type embryos. The founder zebrafish were raised to adult. Following genotype verification using Sanger DNA sequencing, *cav1^+/-^* animals were used for experiments.

### Mouse cornea lymphangiogenesis

VEGFC pellet preparation and implantation were carried out as previously described with minor modifications(*47, 64*). Briefly, pellets containing recombinant VEGFC (5 mg/ml), AIBP (10 mg/ml) in combination with apoA-1 (15 mg/ml), CAV-1 scaffolding domain (CSD; 5 mg/ml), VEGFC with AIBP and ApoA-I or with CSD were prepared by mixing with 10% sucralfate solution (w/v in PBS) and 12% poly-HEMA (w/v in ethanol). The aforementioned components were mixed as following: 5 μl of poly-HEMA, 1 μl of sucralfate, 2 μl VEGFC, 2 μl AIBP, 2 μl apoA-I, 2 μl CSD, and the final volume adjusted to 10 μl with PBS. The mixture droplet was put on the PARAFILM, UV irradiated for 15 minutes, and air-dried in the cell culture hood. Dried pellets were used immediately or stored at 4 °C before usage.

The mouse was anesthetized with isoflurane, and one drop of 0.5% proparacaine HCI was applied to the cornea. Five minutes later, one mouse eye was properly orientated under a dissection microscope. A gentle cut was made with a von Graefe cataract knife from the center of the cornea, and a pocket with ~ 1.5-2 mm^2^ size was generated by inserting the knife toward corneal limbus, with a few gentle waggles inside. The pellets were then inserted into the cornea pocket using forceps and von Graefe cataract knife, with the pellet flattened to secure the implantation. The ophthalmic antibiotic ointment was applied topically to the injured eye. Following 10 days of pellet implantation, the cornea was dissected and washed in PBS for later processing.

### Tissue biopsy

Cutaneous punch biopsy was performed as previously described (*81*). After informed consent was obtained, two contiguous 6 mm full thickness punch biopsy specimens were derived from the medial aspect of the forearm or calf of the affected extremity with an AcuPunch disposable device. Biopsy specimens were immediately place in formalin and processed for subsequent studies.

### Immunohistochemistry (IHC) staining of human lymphedema tissues

AIBP antiserum were generated in rabbits using tag-free recombinant human AIBP proteins. AIBP antibody was subsequently purified from the AIBP antiserum using affinity purification, where AIBP antigen column was prepared by covalently conjugating recombinant AIBP to the amine-reactive, beaded-agarose resin (Thermo Fisher, Pierce NHS-Activated Agarose, Cat # 26196).

The 5um human specimens were deparaffinized, and rehydrated serially through xylene, 100%, 80%, and 70% ethanol, and finally rinsed in milliQ water. The resulting sections were subjected to antigen recovery by immersing in 100°C citrate buffer (pH6.0) for 10 minutes. An incubation with 3% hydrogen peroxide was applied to eliminate the endogenous peroxidase activity. Following rinsing in 1XTBST (diluted from 10X TBST wash buffer, Dako, Cat # S3006), the slides were further washed and blocked in protein block serum-free buffer (Dako, Cat # X0909). The slides were incubated with AIBP antibody for 1 hour at room temperature. Following proper TBST wash, the slides were sequentially incubated with biotinylated anti-rabbit (1:200, Vector, Cat # BA-9500) and Avidin conjugated alkaline phosphatase (Vector, Cat # AK-5000) for 30 minutes at room temperature. AP blue substrate mixture (Vector, Cat # SK-5300) was administered for final color development. The slides were mounted in the aqueous mounting medium.

## Acknowledgments

We thank Dr. Henry Pownall (HMRI) for providing purified human ApoA-I protein. We thank Drs John P. Cooke (HMRI) and Eva Sevick (UT-HSC) for their help and suggestions on this project. This project was supported by grants (NIH HL132155, AHA 16BGIA27790081, AHA 18TPA34250009, and HMRI startup fund) to L.F., AHA (17POST33410671) to X.Y, and AHA (18CDA34110132) to JK, Grants (AHA 19CDA34760260 and a grant from the Oklahoma Center for Adult Stem Cell Research) to P. Y.

## Author contributions

XY and JK designed the experiments, performed most of the experiments and analyzed the data. XY generated the majority of the data in zebrafish and in cell culture models, and JK mainly performed the studies in mice and using human specimens. QG helped on some of the experiment design, experimentation, and data analysis. QY prepared HDL_3_ and recombinant AIBP. PY, JA and PC provided critical suggestions, experimental materials, and technical guidance for this project. SGR provided constructive suggestions and tissues of lymphedema patients and participated in manuscript preparation. XTQ verified some of the results. LF conceived the project, designed experiments, and supervised the studies. XY and LF wrote the manuscript with input from the co-authors.

## Data and materials availability

All data associated with this study are available in the main text or the supplementary materials.

## SUPPLEMENTAL MATERIAL

### Supplemental Methods

#### Zebrafish husbandry

All the wild-type AB and transgenic zebrafish lines were maintained as previously described (*81*) and in accordance with Houston Methodist Research Institute Animal Care and Use Committee (IACUC) regulations and under the appropriate project protocols.

#### MO and mRNA injections

Morpholino antisense oligos (MOs) were synthesized by Gene Tools. The following MOs were used in this study 1) control: 5’-CCTCTTACCTCAGTTACAATTTATA-3’, and 2) *apoa1bp2* (NCBI Gene ID: 557840): 5’-GTGGTTCATCTTGATTTATTCGGC-3’ (*81*). The working concentrations of control and *aopa1bp2* MOs were 0.2 mM and 0.1 mM, respectively. Equal volume of *aopalbp2* MO (0.1 mM) and *aopoalbp2* mRNA (200 ng/μl) were mixed to perform the rescue assays. The working concentration of human *APOA1* mRNA was 100 ng/μl. One nl of MO, mRNA or the combination of MO with mRNA were injected into one-cell stage embryo using the microinjector FemtoJet^®^ 4i (Eppendorf).

#### Chemical treatments

For 4OHT treatment, embryos were treated with 5 μM 4OHT (Sigma, Cat # H7904,) from 6 to 36 hours post fertilization (hfp). For atorvastatin (Sigma, Cat # PZ0001) treatment, embryos were treated with 1 μM atorvastatin from 6 to 120 hpf. All control embryos were incubated with equal amount of ethanol at the indicated time points.

#### Cell culture

Human dermal lymphatic microvascular endothelial cells (hLECs) (Angio-Proteomie, Cat # cAP-0003) were cultured in Endothelial Cell Medium (ScienCell, Cat # 1001). Methyl-β-cyclodextrin (Sigma, Cat # C4555) was prepared in serum-free ECM medium with a working concentration of 10 mM. HDL_3_ isolation was performed as previous described (*81*). The hLECs were pretreated with 200 ng/ml AIBP protein, 100 μg/ml HDL_3_, or the combination of AIBP with HDL_3_ for 4 hours, washed with PBS, and followed by incubation with 100 ng/ml VEGFC (R&D, Cat # 9919) in pre-warmed serum-free ECM medium for 20 min.

#### Lenti-virus infection

The hLECs were transduced with the third generation lentiviral system as previous described (*81*). After the cells were infected with human VEGFR3, VEGFR3^AAA^ or CAV1 overexpression Lenti-virus for 96 hours, cells were subjected to 5 μg/ml puromycin (ThermoFisher, Cat #A1113803)) selection to generate pool cells stably expressing the transduced genes.

#### Quantitative PCR

The qPCR was performed as previously described (*81*). The gene information and the primers used in this study were: murine reference gene *Efla* (GeneBank Accession # NM_010106.2), Ef1a-F (5’-GATCGATCGTCGTTCTGGTAAG-3’) and Ef1a-R (5’-AGTGGAGGGTAGTCAGAGAAG-3’); CD31 (GeneBank Accession # NM_008816.3), CD31-F (5’-CAACAGAGCCAGCAGTATGA) and CD31-R (5’-TGACAACCACCGCAATGA-3’); Lyve1 (GeneBank Accession # NM_053247.4), Lyve1-F (5’-CCTTGTTGGCTGAGACTGTAA) and Lyve1-R (5’-CTAGAGAACACCAGCAACAGTAA-3’); zebrafish reference gene ef1a (GeneBank Accession # NM_131263.1), ef1a-F (5’-ATGCCCTTGATGCCATTCT-3’) and ef1a-R (5’-CCCACAGGTACAGTTCCAATAC-3’); lyve1 (GeneBank Accession # XM_687593.6), lyve1-F (5’-GACAGCTCCCAAACAACAATAAA-3’) and lyve1-R (5’-CTGAGAGGTTGAATGAGAGGAAG-3’); vegfc (Genebank Accession # NM_205734.1), vegfc-F (5’-CGTCTCTTGATGTCTCGGAATG-3’) and vegfc-R (5’-GCTGTTACTTTGGATCCCTCTC-3’); vegfr3 (Genebank Accession # NM_130945.1), vegfr3-F (5’-CAGTGTGGTCACCTGGAATAA-3’) and vegfr3-R (5’-TGGAGCAGTAGAAGCCAATAAA-3’); prox1 (Genebank Accession # NM_131405.2), prox1-F (5’-CGTGATGGATCAAGAGGAAAGA-3’) and prox1-R (5’-CTACCTGGGACATTGCTGTATT-3’); sox18 (Genebank Accession # XM_001337666.3), sox18-F (5’-GGAAGATGTGGGTCTGTCTTC-3’) and sox18-R (5’-TGGGAATGCTGGAGGTTATG-3’); coupTFII (Genebank Accession # NM_131183.1), coupTFII-F (5’-CACAGGTCGCTAACCTATTT-3’) and coupTFII-R (5’-ACAAGGGCTAGTGTACTGAATG-3’).

#### Western blot and Immunoprecipitation

Zebrafish or hLECs were lysed on ice with the RIPA buffer (Pierce) and western blotting was performed as previously described (*81*). Immunoprecipitation was performed using the Pierce Classic Magnetic IP/Co-IP Kit (Thermo Scientific, Cat No. 88804) following the manufacturer’s instruction. The antibodies used were: anti-Phospho-Akt (Ser473) (Cell Signaling, #4060S), anti-AKT (Cell Signaling, #4685S), anti-Phospho-ERK1/2 (Cell Signaling, #4370S), anti-ERK1/2 (Cell Signaling, #9102S), rabbit anti-β-Tubulin (Cell Signaling, #2148S), anti-Phospho-tyrosine (clone 4G10, EMD Millipore, #05-321), anti-Flt4 (Santa Cruz, C20, Cat No. sc321), anti-CAV1 (Cell Signaling, Cat No. 3238S), anti-GAPDH (Cell Signaling, #2118S), anti-mLYVE1 (Angiobio, Cat No. 11034), anti-CD31 (Abcam, Cat No. ab28364), mouse anti-LAMIN A/C (Cell Signaling, Cat No. 4777S), goat anti-Guinea Pig HRP-conjugated antibody (Jackson Labs, Cat No. 106-035-003), goat anti-mouse HRP-conjugated antibody (Jackson Labs, Cat No. 115-035-003), and goat anti-rabbit HRP-conjugated antibody (Jackson Labs, Cat # 111-035-144).

#### Mouse embryonic stem cell (mESC) culture

The mESCs were cultured in feeder- and serum-free environment using ESGRO^®^-2i medium supplemented with leukemia inhibitory factor (LIF), GSK3β inhibitor and MEK1/2 inhibitor (EMD Millipore, Cat #. SF016-100) according to the manufacturer’s instruction. Briefly, T25 flask was coated with 0.1% gelatin solution for 30 min at room temperature, and then 1 × 10^6^ cells were plated onto gelatinized T25 flask in ESGRO^®^-2i medium and incubated at 37 °C with 5% CO_2_. The media were changed daily and mESCs were sub-cultured at a ratio of 1:5 when cells reached 60-90% confluence.

#### Lymphatic endothelial cell (LEC) differentiation

The mESCs differentiation to the endothelial lineage was performed as previously described with the following modifications (*82*). Briefly, mESCs were seeded onto collagen IV-coated plates and maintained in LEC differentiation medium (ESGRO Complete™ basal medium, Cat #. SCR002-500) supplemented with 2 ng/ml BMP4, 10 ng/ml bFGF, 50 ng/ml VEGFA and 50 ng/ml VEGFC for 7 days. Differentiated cells were collected on days 3, 5 and 7 for qPCR, western blot, or FACS analysis.

#### Fluorescence Activated Cell Sorting (FACS) analysis

Cells were dissociated using enzyme-free Hanks’ Balanced Salt Solution (Gibco, Cat No. 13150016), resuspended in 1× DPBS supplemented with 2% heat-inactivated FBS. Cells were then stained with APC anti-CD31 antibody (Bio-legend, Cat No. 102409), Alexa Fluor 488 conjugated LYVE1 Monoclonal Antibody (ALY7), eBioscience™ (Cat No. 53-0443-82) and DAPI, and analyzed by BD LSR II flow cytometer. Data were analyzed using the FlowJo software.

#### VEGFR3 site-direct mutation

VEGFR3 mutation was generated using a PCR-based direct site mutation strategy. Briefly, the plasmid containing wild type VEGFR3 was amplified using primer pair VEGFR3-^AAA^-F (5’-ACGCAGAGTGACGTGGCGTCCGCTGGGGTGCTTCTCGCGGAGATCTTCTCTCTGGGG GCC-3’) and VEGFR3-^AAA^-R (5’-AGAGAGAAGATCTCCGCGAGAAGCACCCCAGCGGACGCCACGTCACTCTGCGTGGT GTAC-3’). The PCR program was 95 °C for 2 min, 18 cycles of 95 °C for 30 s, 68 °C for 1 min per kilobase of plasmid length, then 68 °C for 7 min and hold at 4 °C. One μl DpnI was added to the PCR products to disrupt the plasmid template and incubated at 37 °C for 2 hours, and then 5 μl PCR products were transformed into stbl3 competent cells. The resulting clones were selected, plasmid extracted, and DNA sequenced. The one with expected mutation was used for subsequent experiments.

#### Prox1 staining

Zebrafish larvae at 36 hpf were fixed in 4% paraformaldehyde for 2 hours at room temperature. Following fixation, larvae were washed with PBS, dehydrated through 25%, 50%, 70% and 100% methanol/PBS series and stored at – 20 °C in 100% methanol for later usage. Before usage, larvae were rehydrated through 75%, 50% and 25% methanol/PBS series, washed with PBS/0.1% Tween (PBST), and permeabilized in pre-chilled acetone for 20 min. The resulting animals were then washed with PBST and blocked in 10% BSA/PBST overnight at 4 °C. Following incubation with Prox1 antibody (R&D, Cat No. AF2727) and GFP antibody (ThermoFisher, Cat # A10262) for 3 days at 4 °C, and after proper wash with PBST, the larvae were incubated overnight at 4 °C in dark with secondary antibody diluted in 1% BSA/PBST. Following extensive PBST wash, the immunostained animals were imaged using Olympus FV 3000 confocal microscope.

#### Dissection of the tail skin of neonatal mice

After euthanization, the tails of *Cav1^-/-^* or littermate control at P3 were dissected, with the tail tips cut off. The tail bones and extra connective tissues were then manually removed. Tails were incubated in 20 mM EDTA/PBS overnight at 4°C to facilitate dermis isolation. The dissected dermis was washed with PBS for subsequent immunostaining as described below.

#### Immuno-fluorescence (IF) staining of mouse cornea and tail skin

For IF staining, cornea and tail skin were fixed in 4% paraformaldehyde for 2 hr at room temperature. Subsequently, samples were washed 5 times with 0.05% tween/PBS (0.05% PBS-tw) and blocked with the blocking buffer (1% goat or donkey serum, 1% BSA, and 0.5% Triton X100 in PBS) for 3-4 hr at room temperature. Samples were incubated with LYVE-1 antibody (1:1000; Angiobio, Cat # 11-034) and CD31 antibody (1:1000; BD Biosciences, Cat. 553369) diluted in the blocking solution for 2 days and with secondary antibodies (1:1000; Jackson ImmunoResearch Inc., AB_2338052 and AB_2338372) overnight at 4 °C following proper wash with 0.5% Triton X-100/PBS (0.5% PBS-tx). The immunostained tissues were flat-mounted in the mounting medium (VECTASHIELD, H-1000) and stored at 4 °C before imaging.

#### Confocal imaging and image quantification

Flatmounted tissues were imaged using Nikon A1 confocal microscope. Auto-stitching function was applied to generate the whole cornea image. Z-stacked images were maxi-projected and processed using Image J. Color balance and threshold were adjusted using the same threshold for image quantification.

For quantification of tail lymphatic vessels, images were converted to binary image by applying “Erode” or “Dilate” options as exampled (https://imagej.nih.gov/ij/docs/menus/process.html), and images were skeletonized. Skeleton length was quantified, and pixel value was converted to micrometer (μm).

For quantification of cornea lymphatic vessel (LV) and blood vessel (BV), region of interest (ROI) on each image were chosen based on the location of the VEGFC containing pellets. ROI images were processed using Image J to quantify percentage of LV or BV coverage as was reported(*83*).

#### Data analysis

Statistical comparisons between two groups were analyzed using Student *t* test. One-way ANOVA was used to compare the means of multiple groups. Data were expressed as the mean±SD if n<5 or as the mean±SEM if n≥5. p<0.05 was considered statistically significant. *, p<0.05; **, p<0.01; ***, p<0.001; ****, p<0.0001.

### Supplemental Figures

**Fig. S1.**
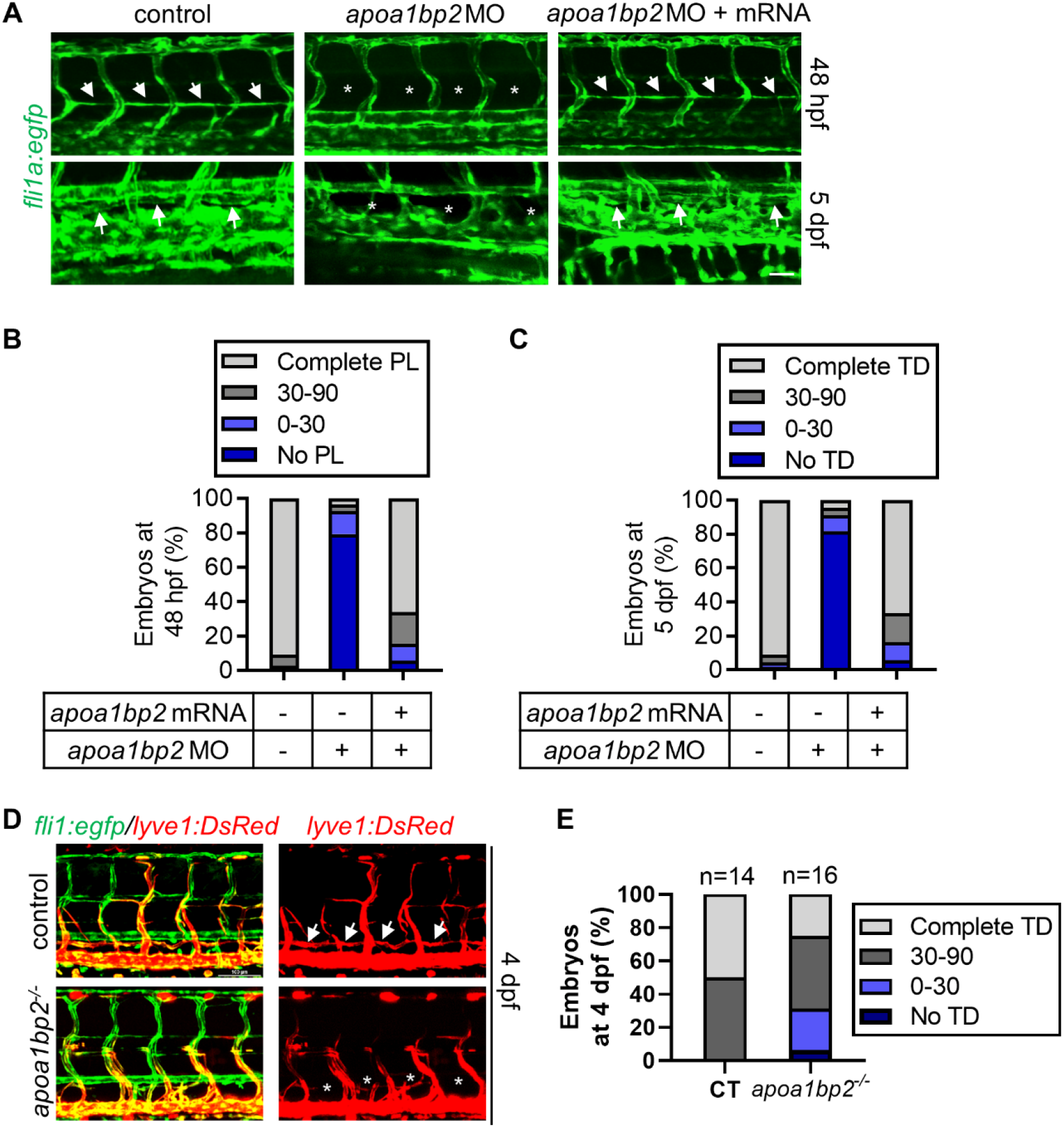
AIBP regulates lymphatic vessel development in zebrafish. **A**, Lymphatic developmental defects in Aibp2 knockdown zebrafish. Confocal images of PL at 48 hpf and TD formation at 5 dpf in the control and Aibp2 null zebrafish. Arrows show PL or TD, and stars denote absent PL or TD. Quantitative data of PL (**B**) and TD formation (**C**) in panel A. n=25 in panel A and =30 in panel C. Scale: 25 μm. **D&E**, TD development in *apoa1bp2^-/-^* and control (CT) *fli1:egpf; lyve1:DsRed* zebrafish (D) and the quantification (E). Scale bar: 100 μm. **, p<0.01.

**Fig. S2.**
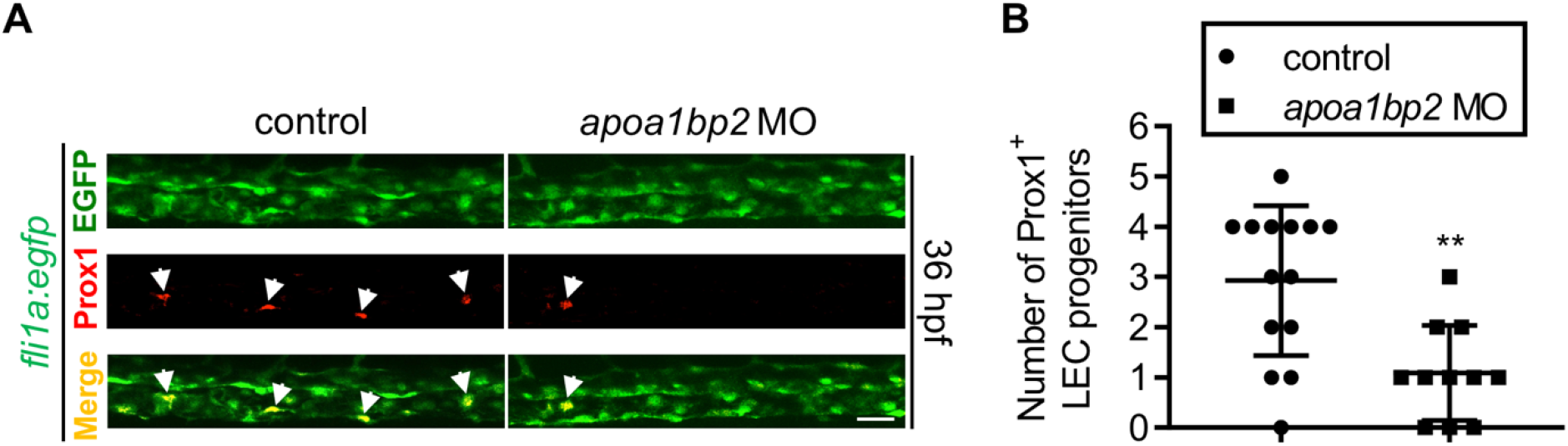
AIBP regulates lymphatic vessel development in zebrafish. (**A**) Aibp2 ablation disrupts LEC specification. Confocal images of LEC progenitors in the CV of control or Aibp2 knockdown zebrafish at 36 hpf. (**B**) Quantitative data of LEC progenitors in panel A. Control or *apoa1bp2* morphants were fixed at 36 hpf, immunostained using Prox1 and EGFP antibodies, and images were captured using confocal microscopy. Arrows indicate LEC progenitors. Scale: 25 μm. **p<0.01.

**Fig. S3.**
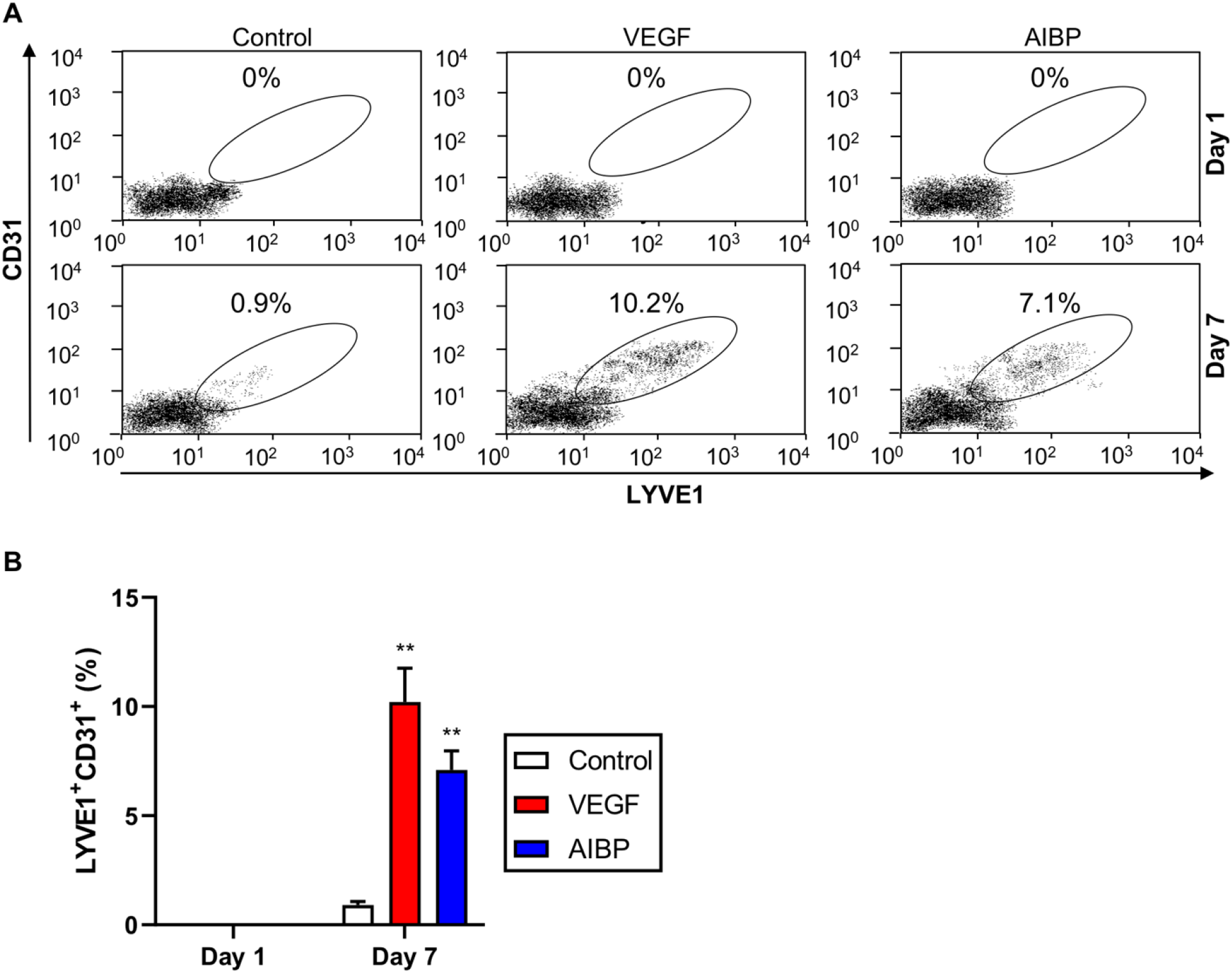
FACS analysis of murine ESC-derived CD31^+^LYVE1^+^ LECs. (**A**) AIBP induces LEC lineage commitment from mESCs. The murine ESCs were subjected to mesoderm and then endothelial differentiation as described in **Fig. 2C**. At day 1 and day 7, the resulting cells were dissociated, immunostained with CD31 and LYVE1 antibodies, fixed with 4% PFA, and used for FACS analysis. The percentage of CD31^+^ and LYVE1^+^ cells were shown. (**B**) Quantitative data of FACS-sorted CD31^+^ and LYVE1^+^ cells in panel (A). **p<0.01.

**Fig. S4.**
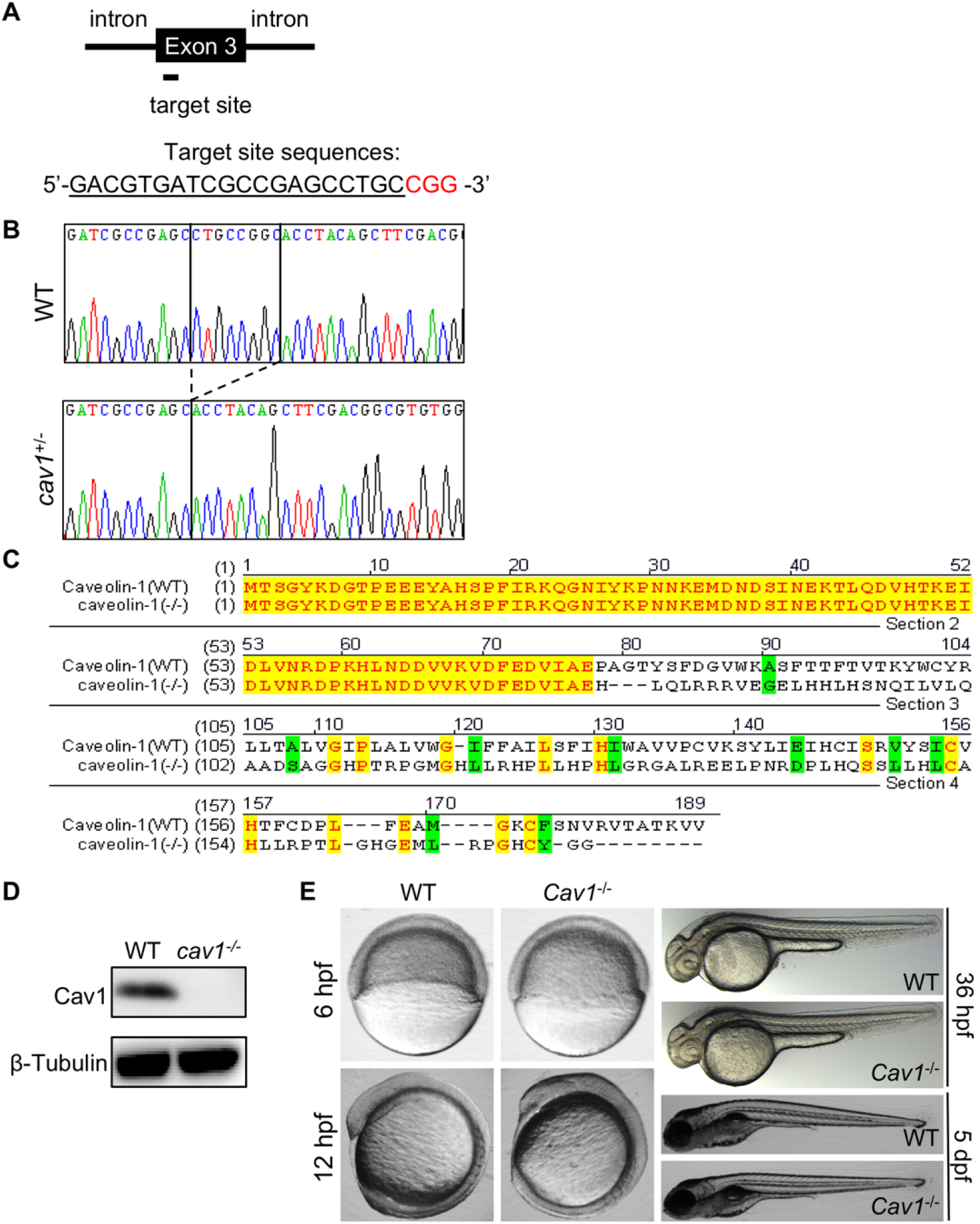
Generation of Cav-1 knockout zebrafish. (**A**) Diagram showing position of the target site and its sequence (underline) in zebrafish *apoa1bp2* locus. PAM sequence (GGG) is shown in red. (**B**) Sanger sequencing result of heterozygous mutants revealed an 8-bp genomic DNA fragment insertion from the target site. The PCR amplicons that span the mutated *apoa1bp2* region were ligated into a T-vector and subsequently transformed into competent cells. Single positive colonies were selected for sequencing. (**C**) The 8-bp insertion resulted in a frame shift that generates a mutated protein. (**D**) Western analysis of pooled 26 hpf zebrafish (n=15) show the absence of Cav-1 expression. (**E**) No gross phenotypic defect observed in Cav-1 knockout zebrafish. Zebrafish embryos at the indicated developmental stages were collected and images of live zebrafish embryos captured. hpf: hour(s) post fertilization. dpf: days post-fertilization. WT: wild type.

**Fig. S5.**
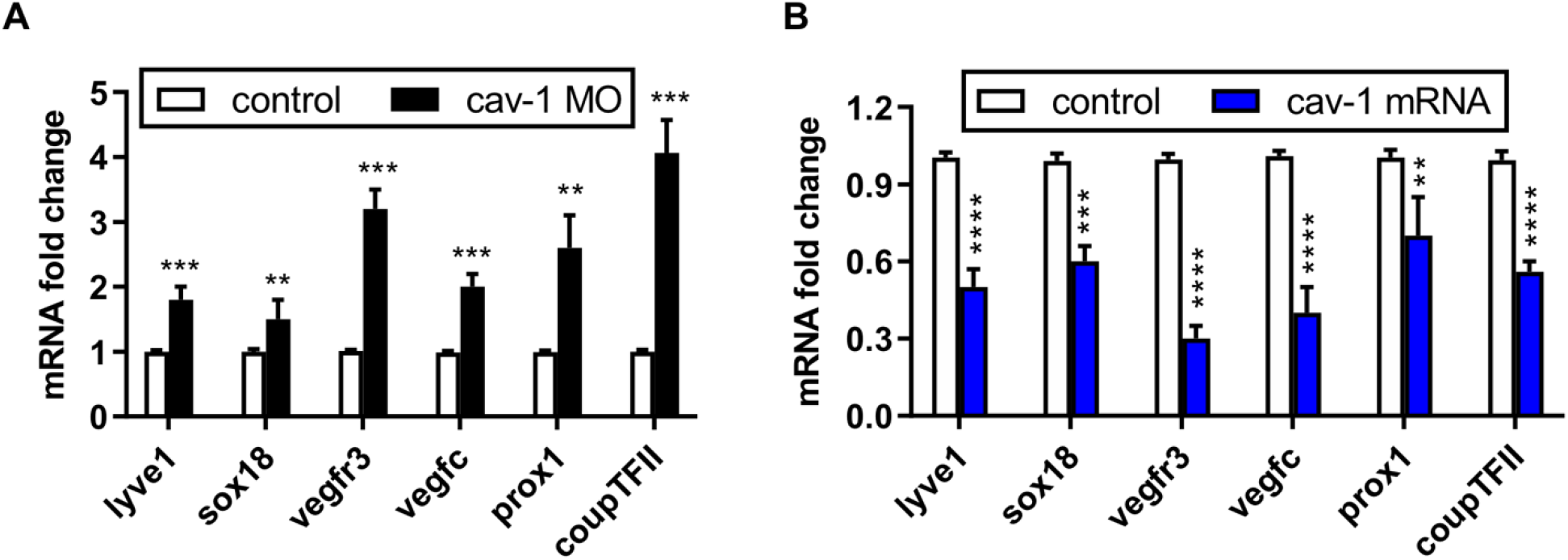
Quantitative PCR analysis of genes regulating lymphatic development in zebrafish. (**A**) Loss of Cav-1 increases LEC gene expression. qPCR analysis of the indicated genes regulating lymphangiogenesis in control and Cav-1 null zebrafish at 96 hpf. (**B**) Cav-1 overexpression reduces LEC gene expression. The zebrafish embryo at one cell stage was injected with Cav-1 mRNA, and the resulting animals or control animals were harvested at 96 hpf, and total RNA extracted for reverse transcription. The indicated genes regulating lymphangiogenesis were analyzed using qPCR in control and Cav-1 overexpressing zebrafish. n=30 per sample. **, p<0.01; ***, p<0.001; ****, p<0.0001.

**Fig. S6.**
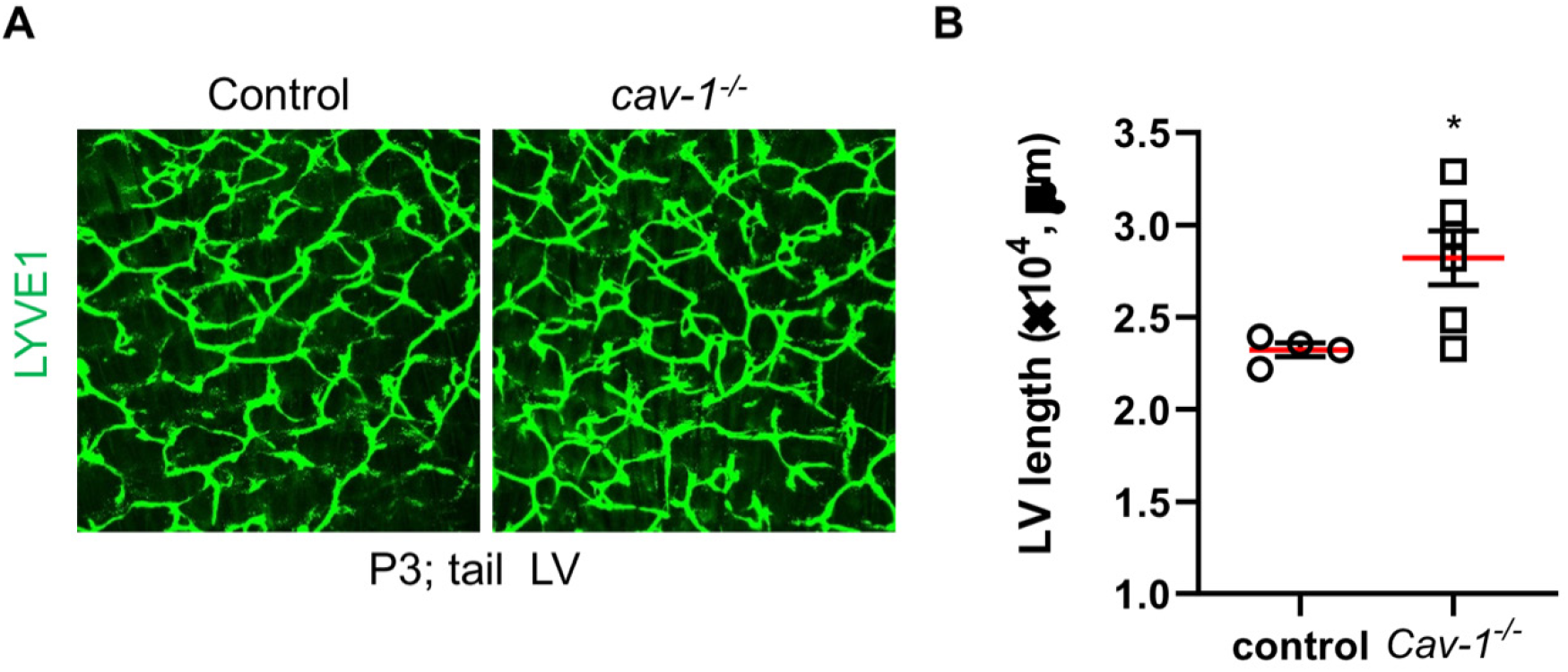
CAV-1 knockout increases tail lymphangiogenesis in neonatal mice. (**A**) The CAV-1 knockout mice were purchased from JAX (stock No. 007083). The tail epidermis of CAV-1 knockout mice and control littermates were dissected from the similar anatomical locations and immunostained using LYVE-1 antibodies. (**B**) Quantification of lymphatic vessel length in panel (A) were performed using ImageJ. *, p<0.05.

**Fig. S7.**
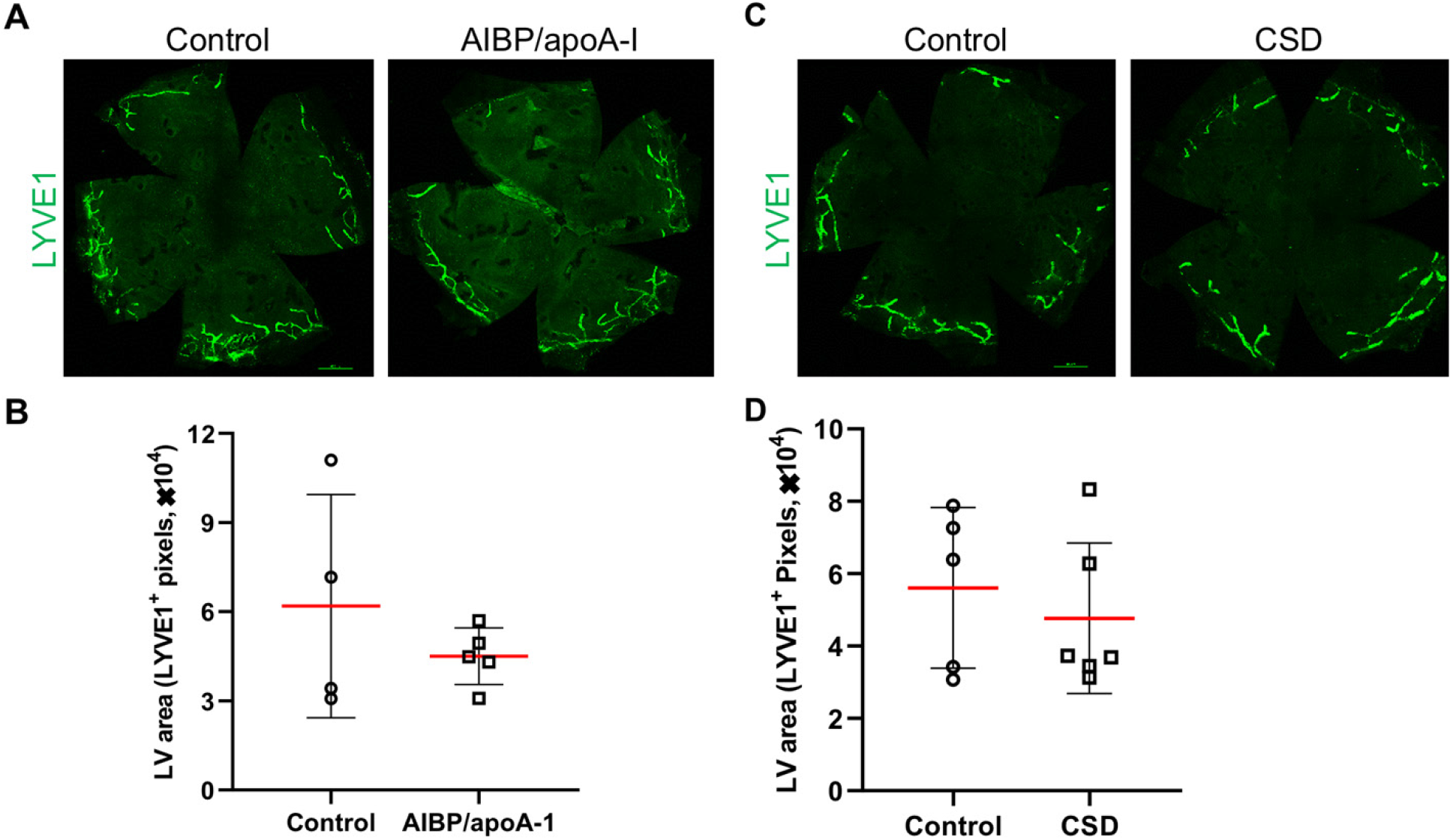
AIBP/apoA-I or CSD per se has no effect on adult lymphangiogenesis. (**A** and **C**), PEG pellets containing recombinant AIBP and apoA-I were prepared and implanted into the corneas of B6 mice, control was implanted with control pellets. (**C** and **D**), Quantification of lymphatic vessel area in panel (A and C).

